# New insights into the extinction and recovery of marine vertebrates across the Permian–Triassic mass extinction event in the Dolomites, Southern Alps, Italy

**DOI:** 10.1101/2025.08.23.671916

**Authors:** Baran Karapunar, Andrzej S. Wolniewicz, Carlo Romano, Péter Ozsvárt, Heriberto Rochín-Bañaga, Evelyn Kustatscher, Stella Z. Buchwald, Francesca Galasso, Donald Davis, Adriana López-Arbarello, Herwig Prinoth, Massimo Bernardi, William J. Foster

**Affiliations:** School of Earth and Environment, University of Leeds, Leeds, LS2 9JT United Kingdom; Department of Earth System Sciences, University of Hamburg, 20146 Hamburg, Germany.; Department of Earth Sciences, University of Cambridge, Cambridge, UK; Institute of Paleobiology, Polish Academy of Sciences, Warsaw, Poland; Paläontologisches Institut und Museum, Universität Zürich, Switzerland; HUN-REN-MTM-ELTE, Research Group for Paleontology, 1431 Budapest, Hungary; Department of Earth Sciences, University of Toronto, Toronto, Ontario M5S 3B1, Canada; Department of Natural History, Tirolean State Museums, Hall in Tirol, Austria; Fundación Miguel Lillo, Unidad Ejecutora Lillo (CONICET-FML), San Miguel de Tucumán, Argentina; Museum Ladin Ciastel de Tor, San Martino in Badia, Italy; MUSE, Museo delle Scienze, Trento, Italy

## Abstract

The impact of the Permian–Triassic mass extinction on marine vertebrates is poorly known due to their scarce fossil record. Here, we report Acrodus sp. [Chondrichthyes], Bobasatrania aff. ladina [Osteichthyes] and an indeterminate actinopterygian from the Changhsingian Bellerophon Formation, and a saurosphargid reptile from the lower Spathian Val Badia Member of the Werfen Formation in the Dolomites, Italy. The saurosphargid is represented by a single vertebra, whose relatively tall neural spine indicates that it might belong to a new species. This is one of the oldest Sauropterygomorpha records worldwide and the first unambiguous occurrence of Saurosphargidae in the western Paleo-Tethys during the Early Triassic. These new findings suggest fishes were more diverse in this paleo-tropical shallow shelf community of the western Paleo-Tethys in the latest Permian than during the Early Triassic. The evolution of the new apex predator niche (Saurosphargidae) relatively shortly after the mass extinction event coincides with the reappearance of predatory cephalopods and precedes the full ecological and taxonomic recovery of benthic groups in the Dolomites community. This contrasts with previous hypotheses about the delayed recovery of higher trophic groups. Furthermore, we dated a surface sample with actinopterygian scales found at a Permian–Triassic boundary outcrop. The U–Pb indicates 248+/-1 Ma as a maximum depositional age, and radiolarian biostratigraphy indicates an Anisian age. This suggests age assignments to surface samples require caution. U–Pb dating further reveals an age discrepancy in the scales and the host rock, likely due to ^206^Pb enrichment in scales during diagenesis.

## INTRODUCTION

The Permian–Triassic mass extinction and the subsequent ecosystem restructuring shifted marine community compositions and the relative diversity of marine clades more than any other event in the Phanerozoic (e.g., Sepkoski 1981; Blois et al. 2013, fig. 1B). Global-scale compilations suggest that marine vertebrates, which predominantly occupy higher trophic levels, underwent lower levels of extinction across the Permian–Triassic transition and rapidly diversified in the Triassic (Scheyer et al. 2014; Romano et al. 2016; Vazquez & Clapham 2017). However, our knowledge of the impact of the event on vertebrates and their post-extinction diversification is more incomplete than for invertebrates, foraminifera, and plants, partly because vertebrate fossils are more scarce (Friedman & Sallan 2012), but also because they remain unreported unless specimens are relatively complete or of particular significance. The scarcity of data hinders assessment of the ecological consequences of the extinction event in local communities across the Permian–Triassic, even in some of the best studied Permian– Triassic successions.

**FIGURE 1.**
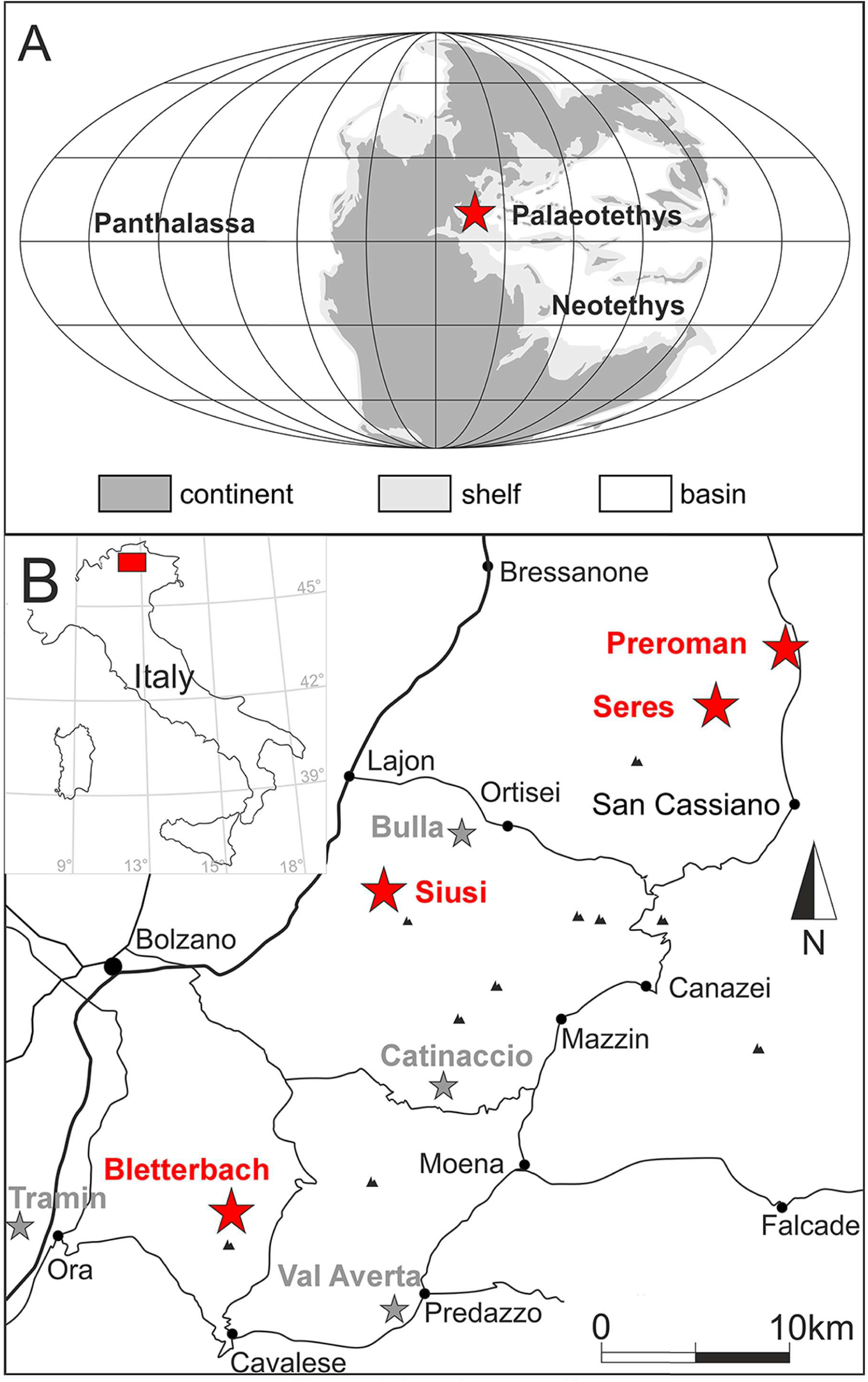
Location maps of the study sites (redrawn after Foster et al. 2017). **A**, Paleogeographic map of the Early Triassic indicating the approximate position of the studied sections in Dolomites, Italy. **B**, Location of the investigated sections in the Dolomites, Italy. The red stars indicate the locations where vertebrates were found: Preroman, Siusi, Seres, Bletterbach section. The grey stars indicate sampled locations without vertebrate findings: Bulla, Catinaccio, Val Averta, and Tramin section. [planned for column width]

The Permian–Triassic transition in the Dolomites, Southern Alps, Italy yields some of the best studied fossiliferous marine successions worldwide, which provide substantial data to analyze the Permian–Triassic mass extinction and subsequent restructuring of marine communities (e.g., Posenato 2009; Hofmann et al. 2015; Foster et al. 2017). The latest Permian– Early Triassic (Changhsingian to Spathian) Bellerophon and Werfen formations in Italy have been studied for nearly two centuries (e.g., Emmrich 1844; Stache 1877, 1878; Ogilvie Gordon 1927; Merla 1930; Leonardi 1935) and have yielded ca. 610 fossil species. The most diverse groups include mollusks (e.g., Wittenburg 1908; Broglio Loriga & Mirabella 1986; Prinoth & Posenato 2007, 2023; Nützel & Karapunar 2023), brachiopods (e.g., Posenato 1998, 2001, 2016), arthropods (e.g., Crasquin et al. 2008), foraminifera and calcareous algae (e.g., Vachard & Krainer 2022), conodonts (e.g., Perri & Farabegoli 2003), and echinoderms (Thompson et al. 2019).

Despite extensive sampling efforts, vertebrate remains, excluding conodonts, are limited to only a few occurrences from the Bellerophon and Werfen formations. Accordi (1955, 1956) reported the actinopterygians *Archaeolepidotus leonardii* (Accordi, 1955), *Bobasatrania ladina* (Accordi, 1955) and *Bobasatrania antiqua* (Accordi, 1955) (=*Paralepidotus*? *moroderi* Accordi, 1955, according to Böttcher 2014) from the latest Changhsingian–Spathian Werfen Formation at Monte Pic, Val Gardena (Dolomites, Italy). Although they were considered Early Triassic by Accordi (1955, 1956), they may be of Permian age (Renato Posenato, pers. comm. in Brinkmann et al. 2010). A single fragment of an *Acrodus*? tooth was reported from the Bellerophon Formation at Mt. Seceda near Ortisei (Italy, Trentino-Alto Adige) by Brandt (2021). Ronchi et al. (2018) reported the trace fossil *Undichna,* interpreted as being produced by a fish swimming, from the Smithian of the Werfen Formation and from the upper Permian (Wuchiapingian) Val Gardena Sandstone (see also Conti 1975, 1977 for fossil fish from the Val Gardena Sandstone). Mostler & Rosner (1984) previously reported a scale of *Gyrolepis*? and teeth belonging to *Saurichthys* and *Colobodus* from the Spathian deposits of the Werfen Formation from the Northern Calcareous Alps, Austria.

Here, we describe newly collected vertebrate remains from the Bellerophon and Werfen formations, including specimens found in horizons just below the extinction boundary (i.e., the formation boundary between the Bellerophon and Werfen formations; Posenato 2019). This new data is then put into context with previous findings to improve our understanding of the impact of the Permian–Triassic mass extinction on marine ecosystems.

## MATERIAL AND METHODS

### Geological setting

The studied sections are located in South Tyrol, northern Italy (Figure 1). These sections record deposition in shallow-shelf environmental settings of the western Paleo-Tethys, at tropical paleolatitudes (e.g., Broglio-Loriga et al. 1990; Massari et al. 1994; Brandner & Keim 2011). The Changhsingian Bellerophon Formation begins with evaporite deposits indicating sabkha environment in a closed basin, and continues with carbonate deposits ranging from marginal marine and restricted low-energy environments to shallow shelf settings (Massari et al. 1994; Posenato 2010). In shallow water sections, the Werfen Formation starts with the uppermost Changhsingian Tesero Member (Farabegoli et al. 2007), which marks the onset of the extinction (Posenato 2019), and continues with low-diversity laminated siltstones and marlstones of the Griesbachian Mazzin Member, which is overlain by the supra-tidal to terrestrial Andraz Member. This is followed by the lower Siusi Member (late Griesbachian), which has the same lithology as the Mazzin Member, but hosts a different species assemblage (Hofmann et al. 2015). The mid-upper Siusi Member records multicolored (dominantly red, brown and green) siliciclastics (Dienerian), the Dienerian reddish Gastropod Oolite Member, the Smithian reddish marlstones and sandstones of the Campil Member; and finally, the grey to yellow colored Spathian Val Badia, Cencenighe and San Lucano members (Broglio Loriga et al. 1988, 1990). The Werfen Formation records repeated transgressive-regressive cycles. The Mazzin, lower Siusi, Val Badia, and lower Campil members represent a subtidal depositional environment; the Andraz, upper Siusi, upper Campil, mid-Cencenighe, and San Lucano members represent a supratidal environment (Broglio Loriga et al. 1983; Hofmann et al. 2015). The mid-Siusi, Gastropod Oolite, Campil and Cencenighe members dominantly represent inner shelf environments.

### Material

Hundreds of surface samples and eight bulk samples (1–3 kg) were collected from the latest Permian–Early Triassic (Changhsingian–Spathian) marine deposits at the following sections: Preroman (46°40’23.3”N 11°54’04.0”E), Siusi (46°32’00.0”N 11°33’40.7”E), Bulla (46°34’13.3”N 11°37’45.4”E), Catinaccio (46°25’25.6”N 11°36’35.6”E), Val Averta (46°17’52.2”N 11°34’18.4”E), Seres (46°38’23.5”N 11°50’25.9”E) and Tramin (46°20’35.1”N 11°14’04.3”E) (Figure 1). These samples yielded invertebrate, vertebrate and trace fossils. The vertebrate fossils were found in the Siusi, Preroman, and Seres sections. No vertebrate fossils were recovered in the samples from Tramin, Catinaccio, Val Averta, or Bulla. In addition to the newly collected samples, previously undescribed vertebrate specimens from the Bellerophon and Werfen formations from the Bletterbach section were located in museum collections (Museum of Nature South Tyrol; Museo delle Scienze di Trento; Geomuseum Radein), and these previously undescribed specimens were also included in this study. The newly collected invertebrate and trace fossils are stored at the Museum Ladin and Museum of Nature South Tyrol, Bolzano, Italy. The studied radiolarian specimens from the Seres section are housed at the Hungarian Natural History Museum. The studied vertebrate fossils are deposited in the Museum of Nature South Tyrol, Bolzano (with the collection number PZO) and Geomuseum Radein, Bolzano (with the collection label GEOM).

### Siusi section

The Siusi (Seis) section (Siegert et al. 2011) is located in a valley carved by the river Rio Bianco (Weissenbach), which has formed several terraces with waterfalls. Fresh exposures of the Bellerophon and Werfen formations are present in cliff faces along the riverbed. The lower part of the section yields root traces indicating a marginal marine setting (Massari et al. 1994). A black shale deposit, approximately 0.5 meters beneath the Bellerophon/Werfen formation boundary, directly underlying nodular limestone, was sampled. A bulk sample of 1.7 kg of black shale was wet-sieved at mesh-sizes of 5 mm, 2 mm, and 0.63 mm. The residues were dried at 40°C and fossils were picked using a binocular microscope. Only three fish teeth were found in the sample: PZO 16551, PZO 16552, and PZO 16553.

### Preroman section

The Preroman section yields the uppermost 50 meters of limestones of the Bellerophon Formation, which corresponds to the Changhsingian (Prinoth & Posenato 2023). This section contains dark colored marine carbonates deposited in a subtidal environment. PZO 16556 was extracted from a dark grey limestone found in a scree deposit derived solely from the Bellerophon Formation limestone.

### Bletterbach section

The succession at the Bletterbach section begins with the early Permian ignimbrites of the Ora Formation and extends up to the Anisian Contrin Formation (Massari et al. 1988; Roghi et al. 2014). GEOM Radein 0035 was found in the Gorz part of the Bletterbach Gorge and originates from the Bellerophon Formation.

The saurosphargid vertebra fossil (PZO 773) originates from the western slope of the Weißhorn, above the Bletterbach Gorge (46°21’27.79’’N 11°26’15.71’’E), at an elevation of 1,945 m above sea level. The horizon of origin belongs to the Val Badia Member of the Werfen Formation, which locally consists of about 40 m of marly limestones interbedded with bioclastic calcarenites containing bivalves, gastropods, and echinoderms. These deposits are often bioturbated and interpreted as deposited in a shelf environment episodically disturbed by storms (Broglio Loriga et al. 1990; Twitchett & Wignall 1996). In the Bletterbach section, the classical invertebrate association diagnostic for this member includes *Tirolites cassianus*, *Scythentolium tirolicum*, *Ladinaticella costata*, and *Werfenella rectecostata* (Neri & Posenato, 1985). The presence of the ammonoid genus *Tirolites* indicates that this member belongs to the lower Spathian (Broglio Loriga et al. 1990; Zhang et al. 2019).

### Seres section

The Seres section (also referred to as the Val Badia or Misci section in the literature; Vachard & Krainer 2022) is composed of strata ranging from the Changhsingian Bellerophon Formation up to the Mazzin Member of the Werfen Formation, which is overlain by the Richthofen Conglomerate (Vachard & Krainer 2022). PZO 16550 was collected from the ostracod-rich facies of the Bellerophon Formation (46°38’23.7”N 11°50’30.4”E), while PZO 16555 was collected from the cropped-out surface of its micritic facies (46°38’23.5”N 11°50’25.9”E).

PZO 16554 was collected from scree close to the formation boundary (46°38’24.0”N 11°50’24.6”E). From this specimen, a sample was cut to prepare a thin section for investigating Radiolaria, and for radiometric dating. The prepared thin section is stored in the Museum of Nature South Tyrol, Bolzano. This thin section revealed that PZO 16554 is a cherty limestone with radiolarians and ostracods. To extract radiolarians, a cut part of this sample (PZO 16554) was processed using a standard dissolution method. It was first placed in approximately 10– 12% HCl for one hour. The residues were then washed successively through 250, 125, and 63 μm sieves and then dried. Next, the insoluble chert residue was dried out after dissolution of the sample and then placed in 5% hydrofluoric acid for 24 hours. This procedure was repeated four times for each sample.

### U–Pb method

U–Pb isotopic analyses on samples from PZO 16554 were conducted at the University of Toronto using an Agilent 7900 ICPMS and an NWR193 excimer laser system. U–Pb data (^238^U, ^206^Pb, and ^207^Pb) were collected using scan-lines over the surface of the host rock and fish scales, resulting in hundreds of cycle data per line (Hoareau et al., 2021; Davis & Rochín-Bañaga, 2021).

Each scan-line analysis, 5 mm long, was pre-ablated at a fast scan rate, 200 μm/sec, and high frequency, 20 Hz, using a larger diameter beam to remove surface contamination. U–Pb line analyses were conducted with a laser wavelength of 193 nm, fluence of about 4 J/cm^2^, diameter beam of 120 µm, and frequency of 10 Hz at a rate of 15 μm/sec. Baselines were accumulated on each scan-line for 20 seconds prior to opening the laser. LA–ICPMS parameters for U–Pb analyses are found in Supplementary Material (Table S1).

The glass standard NIST612, Madagascar apatite standard (MAD2), and Walnut Canyon standard (WC-1) were used to correct for mass and oxide elemental fractionation bias. The WC-1 was used as a matrix-matched reference for the limestone, whereas the MAD2 was used as a matrix-matched reference for the fish scales. The uncertainty of our U–Pb ages includes measurement errors in mass signals from the sample, NIST612, MAD2 and WC-1 standard measurements. The age of the limestone host rock has been corrected by a factor of 1.07 based on comparing the measured age of WC-1 to its true age as determined by ID-TIMS (Roberts et al., 2017). An additional calcite standard (TARIM), of 208.5 ± 0.6 Ma (Zhang et al., 2023) was measured to provide control on accuracy and precision. The age of TARIM was measured here at 207 ± 2 Ma (2σ, MSWD = 0.65) with a fixed ^207^Pb/^206^Pb ratio of 0.86. No age correction was applied for the fish scales. An additional standard, Madagascar apatite standard (MAD1), was measured to provide control of accuracy and precision for the fish scales. The ages of the MAD2 and MAD1 apatite were previously measured by ID-TIMS at 474.3 ± 0.4 Ma and 486.6 ± 0.9 Ma, respectively, by Thomson et al. (2012).

#### U–Pb Data Processing

Data were processed as single measurement cycles to exploit the maximum U over common Pb heterogenicity in the samples and reveal the maximum spread along the mixing line. U–Pb regression was carried out using a Bayesian statistical program (UtilChron) that regresses mass-count data in a 3D signal space (Davis & Rochín-Bañaga 2021). U–Pb data are plotted and regressed using Tera-Wasserburg (T-W) concordia diagrams.

### Photography

PZO 16554 and PZO 16555 were photographed with a Nikon D7000 digital camera equipped with a Nikon AF-S micro 60 mm objective under white and ultraviolet (UV) light (Krantz UV Lamp I 361). PZO 773 was photographed using an Olympus XZ-1 digital camera under white light. Scanning Electron Microscope (SEM) imaging of radiolarians was performed using a ThermoFisher Phenom XL desktop electron microscope equipped with a CeB6 electron source. Measurement conditions were 10–15 kV voltage with a 5.3 nA sample current in 60 Pa low vacuum mode. SEM imaging of fish teeth was performed using a Hitachi TM4000Plus tabletop microscope at 15 kV acceleration voltage under standard vacuum mode. SEM imaging was conducted without coating the specimens. For microscope imaging of PZO 16556, the specimen was coated with ammonium chloride and photographed using a Leica DFC 480 digital camera connected to a Leica M165 FC Stereo Microscope.

### Taxonomy

The terminology for elasmobranch tooth morphology follows Romano et al. (2019, fig. 1), which is based on Fischer et al. (2010). For the morphology of the tooth plate we follow Böttcher (2014). Open nomenclature is applied according to the conventions of Bengtson (1988).

> SYSTEMATIC PALAEONTOLOGY
>
> Clade EUTELEOSTOMI Nelson, 1994
>
> Class CHONDRICHTHYES Huxley, 1880
>
> [by C. Romano]
>
> Cohort EUSELACHII Hay, 1902
>
> Order HYBODONTIFORMES Maisey, 1975
>
> Family ACRODONTIDAE Casier, 1959
>
> Genus *ACRODUS* Agassiz, 1843
>
> *ACRODUS* sp.
>
> Figure 2

**FIGURE 2.**
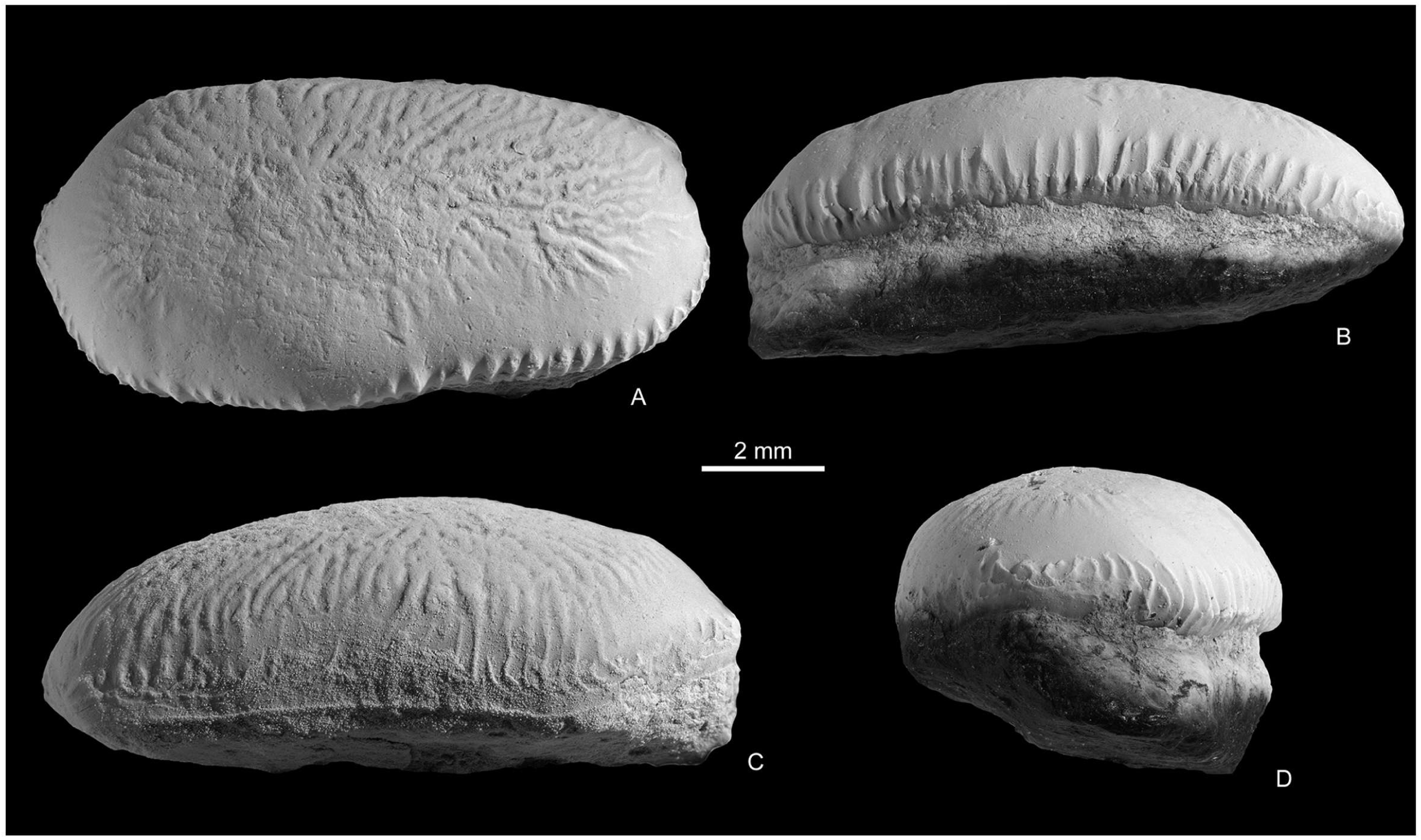
*Acrodus* sp., a single tooth with complete crown and root, PZO 16556, from the Changhsingian Bellerophon Formation, at Preroman section. **A**, occlusal view; **B**, lingual view; **C**, labial view; **D**, mesial? view. [planned for page width]

**Referred specimen–** PZO 16556, a single tooth with complete crown and root. The crown is partly worn.

**Horizon, locality, and age–** Changhsingian, Bellerophon Formation, Preroman section.

**Description–** The tooth has a mesio-distal length of ca. 10.7 mm, a labio-lingual width of about 5.2 mm, and a maximum apico-basal height of ca. 4.3 mm.

The crown has a convex cross-section in both mesio-distal and labio-lingual direction. It lacks any cusps or cusplets. The crown juts out beyond the root on all sides. It is somewhat oval in occlusal view, being longer than wide, with one end (probably the mesial) being pointed and the other one (probably the distal) rounded. While its labial side is rather straight in occlusal view, its lingual face is gently convex just anterior (mesial) and posterior (distal) to the middle of the tooth. A labial protuberance is missing. Ornamentation of the crown consists of reticulated striae, which radiate from the centrally located apex of the crown towards all sides. The striae are, however, weak due to the fact that part of the crown is worn, with the mesial, lingual and distal slopes being smooth. The striae are more pronounced and vertically arranged on the lingual side, near the crown-root junction. An occlusal crest is not discernible, which could be due to the wear.

The root is relatively low; on the lingual face it is apico-basally slightly less deep than the crown. Labially, the root shows a distinct depression. The labial depression reaches the mesial and distal end of the tooth. A sulcus follows the crown-root junction on the lingual side of the tooth. The base of the root is flat. Foramina are absent.

**Remarks–** The morphology of the tooth agrees well with that of flat-crowned, durophagous hybodontiforms, to which it is compared below. The reticulated crown ornamentation resembles that of the Middle Triassic *Palaeobates angustissimus*. However, referral of the tooth to the Triassic *Palaeobates* is hindered due to the relatively low root and also the apparent lack of an occlusal crest (Cappetta 2012). A close relationship with *Asteracanthus* is also unlikely for the same reasons. The tooth morphology is consistent with the anterior teeth in species of *Acrodus*. A major difference is, however, that in *Acrodus*’ teeth the labial contour of the crown is convex (straight in PZO 16556) and the lingual contour is concave (convex in PZO 16556). Tooth histology has not been studied herein due to the fact that it is the only elasmobranch tooth from this section known so far. While the crown of *Palaeobates* contains orthodentine, that of *Acrodus* is made of osteodentine (e.g., Cappetta 2012; Romano & Brinkmann 2010).

Originating from the uppermost Permian strata, the PZO 16556 represents one of the few Paleozoic occurrences of *Acrodus*, a primarily Mesozoic genus (see Discussion below). The present study confirms the presence of this genus prior to the Permian–Triassic mass extinction event.

> Class OSTEICHTHYES Huxley, 1880
>
> [by C. Romano]
>
> Subclass ACTINOPTERYGII Goodrich, 1930
>
> Family BOBASATRANIIDAE Stensiö, 1932
>
> Genus *BOBASATRANIA* White, 1932
>
> *BOBASATRANIA* aff. *LADINA* (Accordi, 1955)
>
> Figure 3

**FIGURE 3.**
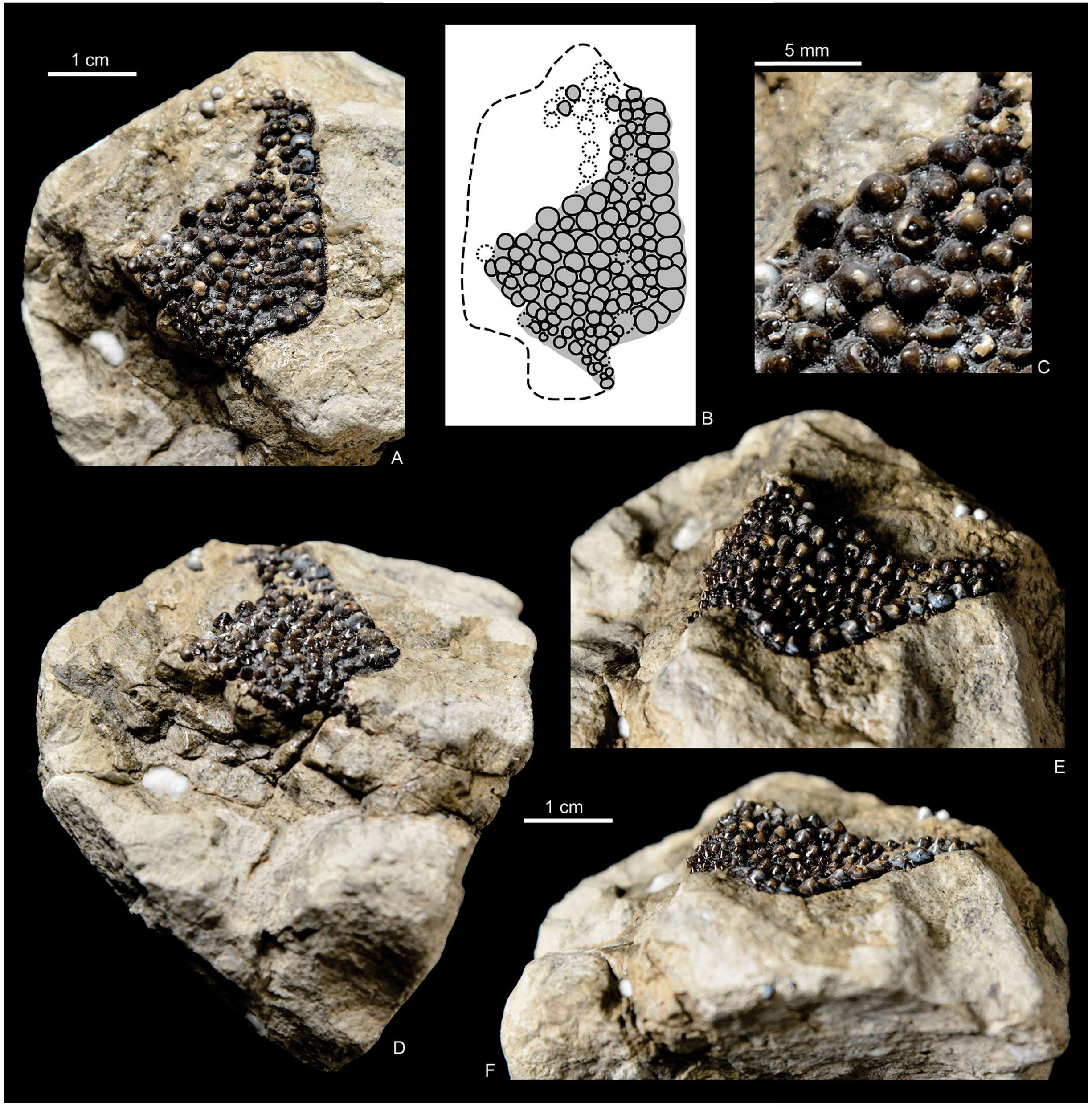
*Bobasatrania* aff. *ladina* (Accordi, 1955), an isolated fragment of a phyllodont tooth plate covered with numerous, close-set low-conical teeth, GEOM Radein 0035, from the Changhsingian Bellerophon Formation at the Bletterbach section. **A**, occlusal view; **B**, Drawing of the tooth plate in occlusal view (from A); **C**, close-up of tooth-plate, showing the superimposed teeth; **D**, oblique posterior-occlusal view; **E**, oblique occlusal-lateral view; **F**, lateral view. [planned for page width]

aff. 1955 *Paralepidotus? ladinus.* n. sp. Accordi, p. 21, pl. 2, pl. 3, figs 1, 5.

**Referred specimen–** GEOM Radein 0035, an isolated fragment of a phyllodont tooth plate covered with numerous, close-set low-conical teeth.

**Horizon, locality, and age–** GEOM Radein 0035 was recovered from the Changhsingian Bellerophon Formation at the Bletterbach section.

**Description–** The tooth plate fragment has a preserved length of ca. 35 mm and a maximal width of about 24 mm. It is presumed that most of the bone is preserved and therefore its original size was only slightly larger. The fragment is bound by an intact lateral margin, which is mostly straight but bends medially at both ends, forming obtuse, but nearly rectangular angles. At both the anterior and posterior ends of the tooth plate, the bone forms processi, of which the presumed posterior one is more complete. A series of large, shallow conical teeth follows the straight lateral margin. The medially neighboring teeth are of the same shape but distinctly smaller. The teeth become slightly larger again towards the center of the bone, near the fracture. On the opposite lateral side, the teeth become smaller again. The oral surface of the dental plate is concave in the areas where the small teeth are situated, whereas it is distinctly convex in the area of the medially located larger teeth.

On the break surface, a cross-section of the tooth plate is exposed. At least two layers of teeth are observed (Figure 3C). These superimposed layers of replacement teeth are typical for phyllodont tooth plates (see e.g., Böttcher 2014).

**Remarks–** The morphology of the phyllodont tooth plate under study agrees well with that of the deep-bodied ray-finned fish *Bobasatrania*. *Bobasatrania* is a typical component of Early Triassic ichthyofaunas; this genus reached a nearly global distribution during this epoch (e.g., Tintori et al. 2014). It has its first occurrence in the latest Permian and survived until the Middle Triassic (Böttcher 2014). Its Permian record is primarily based on isolated dental plates from northern Italy (Dolomites) and East Greenland (Stemmerik et al. 2001; Tintori et al. 2014; Surlyk et al. 2017). From the latest Permian of the Dolomites, Accordi (1955) described two disarticulated tooth plates as *Paralepidotus*? *antiquus* and *P.*? *moroderi*, respectively, and from the Early Triassic of the same region a tooth plate as *P.*? *ladinus*. Böttcher (2014) reassigned *P.*? *antiquus*, *P.*? *ladinus* and *P.*? *moroderi* to *Bobasatrania*. Furthermore Böttcher (2014) considered *Bobasatrania antiqua* and *B. moroderi* to be likely conspecific. He also suggested that the tooth plate of “*P. moroderi*” figured by Pozzi (1993: fig. 45) should rather be referred to as *B. ladina*. We figure the holotype of *Bobasatrania ladina* (Accordi, 1955) for comparison with our material (Figure 4).

**FIGURE 4.**
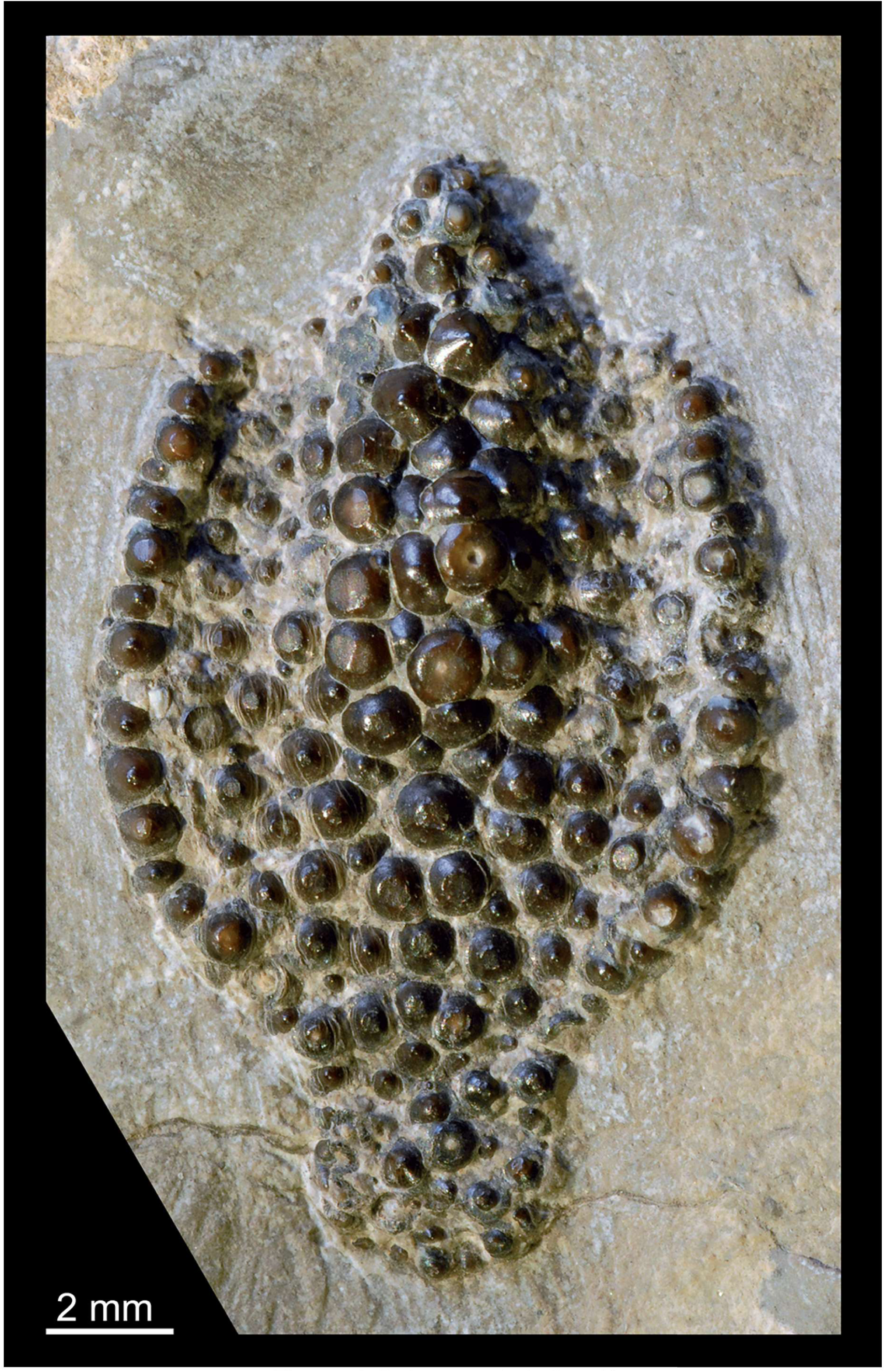
*Bobasatrania ladina* (Accordi, 1955), holotype, tooth plate in occlusal view, from the Monte Pic, Dolomites, stored in the Museum Gherdëina. [planned for column width]

Although the preservation of the tooth plate fragment from the Bletterbach section is imperfect, one of its lateral margins is mostly complete and shows distinct angles at its anterior and posterior ends, both being slightly larger than 90°. The mostly straight lateral margin is seamed with a row of large conical teeth, which are bordered medially by smaller ones, with larger teeth again occurring along the plate’s central midline. This pattern of tooth distribution is similar to the type specimen of *Bobasatrania ladina* (cf. Accordi 1955: pl. 2, figure on top; Figure 4), and it suggests that the fragment shows the postero-lateral portion of the tooth plate. However, the postero-lateral angle in the lateral margin is clearly more acute in the specimen studied herein, and the lateral border is convex in the holotype but straight in GEOM Radein 0035, preventing its attribution to *B. ladina*. The distribution of large and small teeth in *B. antiqua* and *B. moroderi* is different to the specimen studied herein as there does not seem to be a continuous row of larger teeth along the edges in these two species (Accordi 1955: pl. 2, figure at the bottom, pl. 3., figs 2, 4). Based on the comparisons above and in light of the current knowledge, the Bletterbach tooth plate cannot be decidedly attributed at the species level, but because of some similarities with *B. ladina* we refer to it as *Bobasatrania* aff. *ladina*.

Based on the shape of the preserved part of the dental plate fragment, we infer a somewhat leaf-shaped outline of the tooth plate, somewhat broader than the holotype of *B. ladina*. Regarding its outline and due to the curvature of the oral face, which exhibits a central and two lateral ridges with larger teeth separated by two furrows with smaller teeth, the element in question most likely corresponds to an upper tooth plate (cf. Böttcher 2014).

> BOBASATRANIA sp.
>
> Figure 5

**FIGURE 5.**
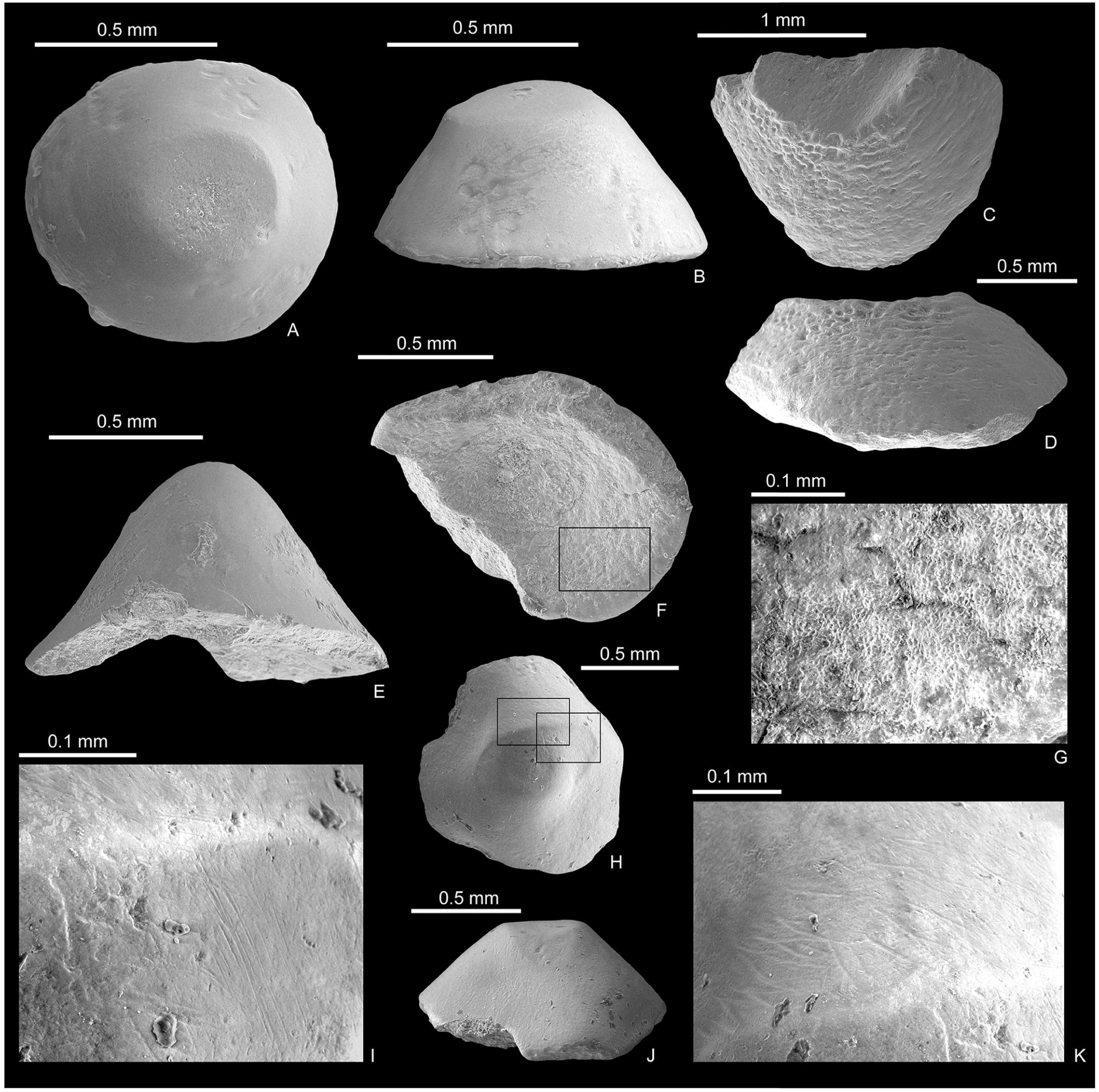
*Bobasatrania* sp. single tooth elements from the Changhsingian Bellerophon Formation at Seres and Siusi sections. **A–B,** PZO 16552, Siusi section, 20 cm below the Tesero Member, **A,** occlusal view; **B,** lateral view. **C–D,** PZO 16553, Siusi section, 20 cm below the Tesero Member, **C,** occlusal view; **D,** lateral view. **E–G,** PZO 16551, Siusi section, 20 cm below the Tesero Member, **E,** lateral view; **F,** basal view; **G,** close up of basal view (rectangle in F) showing dentine tubules (or lacunae?) and possibly mineralized collagen fibres in the bone fracture surface. **H–K,** PZO 16550, Seres section, **H,** occlusal view; **I**, close-up of occlusal view (upper left rectangle in H), showing scratch marks; **J**, lateral view; **K**, close-up of occlusal view (lower right rectangle in H), showing scratch marks on tooth surface. [planned for page width]

**Referred specimens–** PZO 16550, a single tooth element from the Changhsingian Bellerophon Formation of the Seres section. PZO 16551, PZO 16552, PZO 16553, three single tooth specimens from the Changhsingian Bellerophon Formation at the Siusi section, 20 cm below the Tesero Member.

**Horizon, locality, and age–** The material is from the Bellerophon Formation of the Seres and Siusi sections.

**Description–** The teeth are all flat-conical with a rounded apex. Some teeth show signs of wear on their surface.

**Remarks–** The teeth resemble phyllodont teeth (mainly acrodin caps) of tooth plates of *Bobasatrania* (cf. Böttcher 2014), but we refrain from assigning them at species level. At their bases, the teeth contain remains of dentine and probably the dentine tubules are in part visible as openings (Figure 5G). The specimen figured in Vachard & Krainer (2022, pl. 25, fig. 14) from the Seres section is also herein considered as belonging to *Bobasatrania*.

> Order PTYCHOLEPIFORMES Andrews et al., 1967
>
> Family PTYCHOLEPIDAE Brough, 1939
>
> PTYCHOLEPIDAE gen. et sp. indet.
>
> Figure 6

**FIGURE 6.**
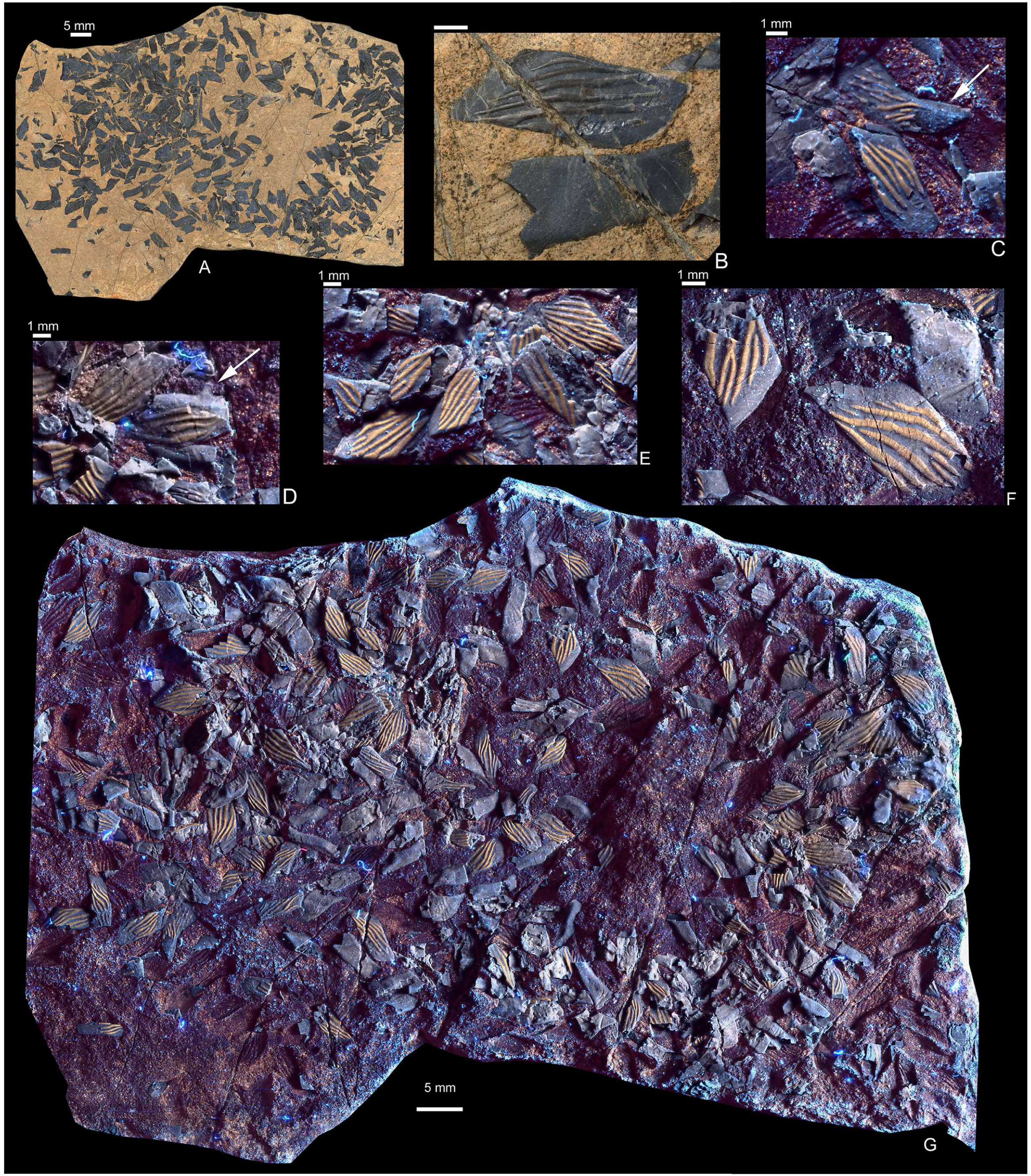
Ptycholepidae gen. et sp. indet., PZO 16554, a rock slab with scattered disarticulated and semi-articulated rhomboid scales and other elements (e.g. lepidotrichia elements), Anisian–Ladinian (Middle Triassic) Buchenstein Formation (Illyrian according to radiolarians, Olenekian–Anisian transition according to U–Pb dating). **A**, The rock slab with scales under white light; **B**, close-up of two scales under white light in lateral (above) and medial (below) aspect. **C**, close-up of scales under UV light, showing ganoine layer only on the ridges (appear in yellow colour), arrow pointing at antero-dorsal process; **D**, scales under UV light, arrow pointing at small dorsal peg; **E–F**, scales under UV light, showing ganoine layer only on the ridges (appear in yellow colour), with some bifurcations of the ridges; **G,** The rock slab with scales under UV light, showing ganoine layer only on the ridges (appear in yellow colour). [planned for page width]

**Referred specimen–** PZO 16554, a single block with scattered disarticulated rhomboid scales and other elements (e.g. lepidotrichia segments). Some of the scales are damaged and therefore incomplete, while others are only preserved as impressions.

**Horizon, locality, and age–** The specimen was collected in the Seres section on the Permian/Triassic boundary outcrop, but based on our analyses it originates from the Anisian– Ladinian (Middle Triassic) Buchenstein Formation (Illyrian [latest Anisian] according to radiolarians, latest Olenekian–Aegean [earliest Anisian] according to U–Pb dating; see below). It was seemingly carried down from the overlying Buchenstein beds outcropping at a distant location.

**Description–** The semi-articulated scales are rhomboid in outline and distinctly longer than deep. The largest scales measure about 6 mm in length and 3.5−4 mm in depth. A few scales show a small dorsal peg (Figure 6D), indicating peg-and-socket articulation, but most do not have such a process. Their anterodorsal and posteroventral portions of the scales are slightly elongated, with the scale’s margins meeting in an acute angle. A distinct antero-dorsal process can only be observed in some scales (Figure 6C). A relatively wide area along the anterior scale margin is smooth; this area was in vivo overlapped by the preceding scale. The free field is ornamented with distinct ganoine ridges, which run from anterodorsal to posteroventral. The ridges are gently curved and mostly separate – they anastomose only on some of the scales, and usually not more than once, while in most scales they do not. Denticulation of the posterior scale margin has not been observed. Some thin, elongate elements scattered among the scales are interpreted as fin ray segments or potentially radials.

**Remarks–** The fact that all scales are elongate-rhombic, being usually much longer than deep, and because of their ornamentation consisting of distinct obliquely antero-posteriorly oriented ganoine ridges suggests affinity with the Mesozoic family Ptycholepidae (cf. Brough 1939, Schaeffer et al. 1975). Referral to *Gyrolepis* is excluded because its scales typically have more numerous but less pronounced ridges on the free field (Bürgin 1992), and a similar condition is seen in scales of *Pteronisculus* (e.g. Romano et al. 2019). In ptycholepids, ganoine is typically restricted to the ridges (Nielsen 1942, Schaeffer et al. 1975, Bürgin 1992). Although the posterior border of the scales is usually denticulated in ptycholepids (e.g., Nielsen 1942), this feature can also be absent in some scales (Brough 1939).

Ptycholepidae includes six genera: *Acrorhabdus*, *Ardoreosomus*, *Boreosomus*, *Chungkingichthys*, *Ptycholepis* and *Youchoulepis*. These taxa have relatively similar scale coverage, except that in *Ardoreosomus*, *Boreosomus* and *Chungkingichthys* the flank scales are equally deep and long, with those located in a more dorsal and ventral position being less deep than long, whereas in *Ptycholepis* and *Youchoulepis* all scales are longer than deep (Schaeffer et al. 1975, Bürgin 1992, Romano et al. 2019). Therefore, based on scale shape alone, PZO 16554 is more similar to *Ardoreosomus*, *Boreosomus* and *Chungkingichthys* than to *Ptycholepis* and *Youchoulepis.* In *Ardoreosomus*, however, the ridges are less pronounced compared to PZO 16554. Due to the imperfect preservation and especially in the absence of cranial features, we refrain from a taxonomic assignment below family level.

> ACTINOPTERYGII gen. et sp. indet.
>
> Figure 7

**FIGURE 7.**
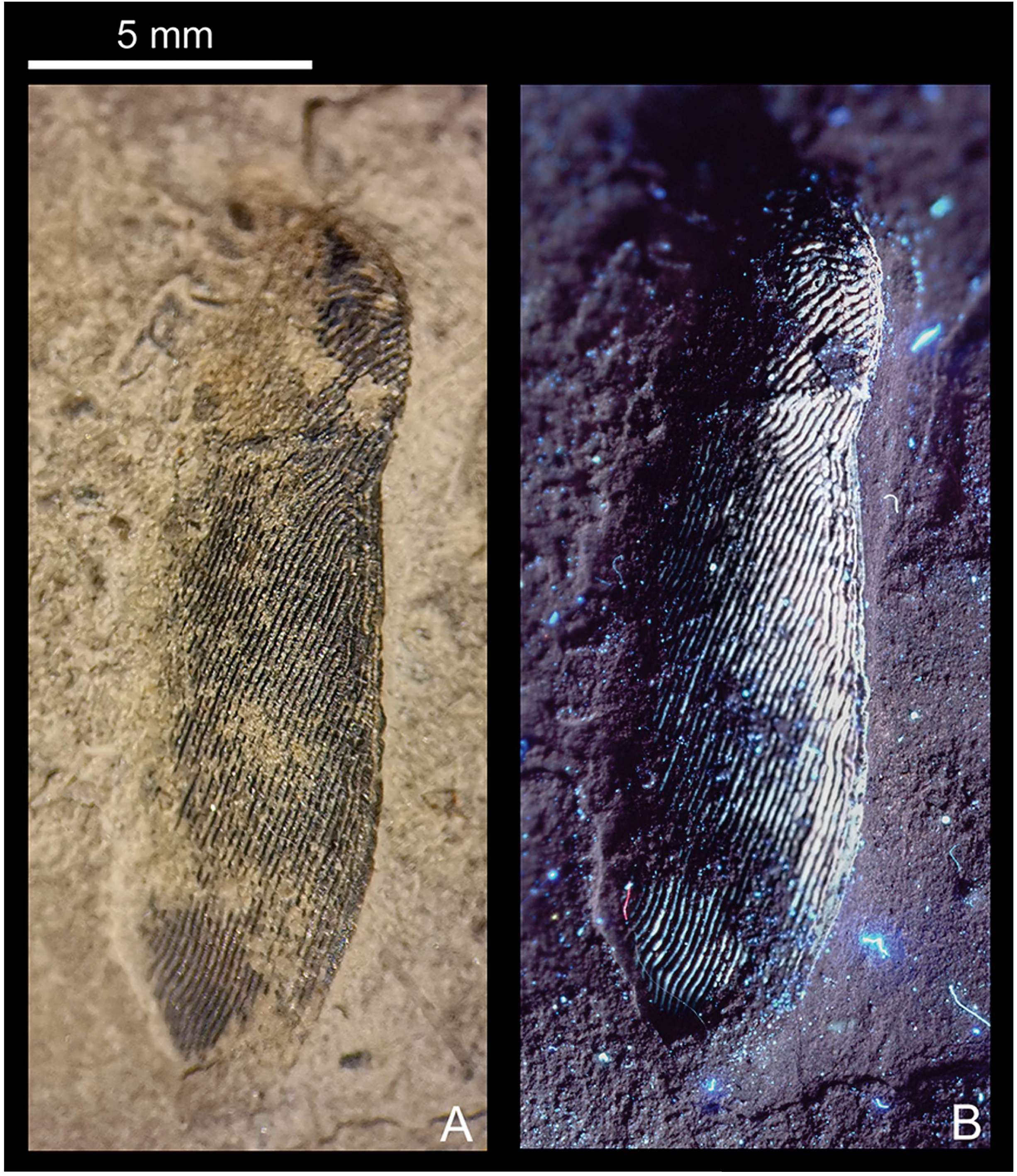
Actinopterygii gen. et sp. indet., PZO 16555, single, isolated bone from the Changhsingian Bellerophon Formation at the Seres section. **A**, the specimen under white light; **B**, the specimen under UV light. [planned for column width]

**Referred specimen–** PZO 16555, an isolated plate-like bone.

**Horizon, locality, and age–** An isolated element from the Changhsingian Bellerophon Formation collected at the Seres section.

**Description–** The plate-like bone is elongate and ovoid in outline. It is ca. 15 mm long. It bears a slight constriction near one of its ends. Its greatest width (4 mm) is observed near its opposite end. At the level of the constriction, it is about 3 mm wide. The external surface of this element is notably ornamented with striae, which run obliquely, nearly parallel to the longitudinal axis of the bone. The striae are mostly straight but gently curved near the constricted region. A sensory canal traverses the bone almost perpendicularly to its longitudinal axis near the constriction.

**Remarks–** Based on the overall shape of the bone, its ornamentation pattern and the location and course of the sensory canal, it is hypothesized here that the element in question is either a deepened flank scale or, possibly, a supracleithrum, with the sensory canal corresponding to a section of the lateral line canal in either case.

Deepened flank scales occur in several Triassic taxa (e.g., Mutter & Herzog 2004), but they are typically of a rhomboid shape, hence different to the outline of PZO 16555. Deepened flank scales with a rounded contour are present in some species of *Saurichthys*, such as *Saurichthys* aff. *dayi* (mid-lateral scales, Kogan 2011). These scales are also ornamented with subvertical striae, but only ventral to the lateral line canal, while dorsally to it, the ornamentation consists of horizontally aligned tubercles (also in *S. madagascariensis*, Kogan & Romano 2016) − in contrast to PZO 16555. Although a vertical striation is also observed on deepened scales of *Bobasatrania*, the outline of these scales is typically rhomboid and no constriction is observed at the level of the lateral line canal. Moreover, no dorsal peg is present (Stensiö 1921, 1932).

Considering the element in question as a supracleithrum, no similarities could be found with known taxa. Affinity with *Bobasatrania* or ptycholepids is doubtful due to differences in the outline of their supracleithra and course of the sensory canal through this bone (cf. Stensiö 1932; Bürgin 1992). The supracleithrum of *Archaeolepidotus* from Val Gardena (northern Italy) has not been described by Accordi (1955, 1956). We herein illustrate the type specimen of *Archaeolepidotus leonardii* Accordi, 1955 (Figure 8), the supracleithrum of which cannot be observed due to preservation. *Archaeolepidotus* was presumed to come from the Lower Triassic Werfen Formation (see Introduction), but might be from the Upper Permian Bellerophon Formation (Renato Posenato, pers. comm. in Brinkmann et al. 2010). In PZO 16555, the sensory line transverses the bone almost perpendicular to its longitudinal axis, which is unusual for a supracleithrum.

**FIGURE 8.**
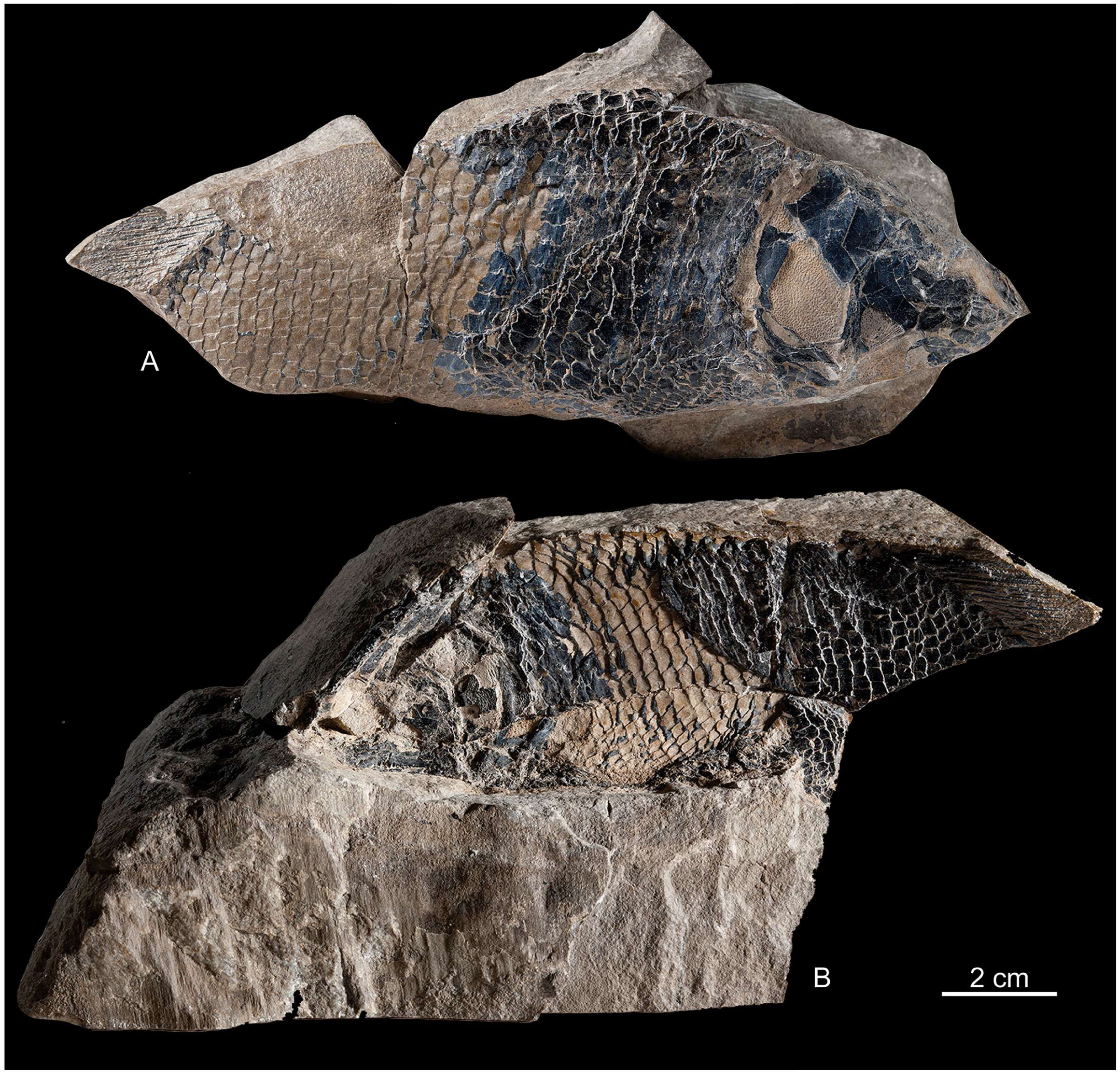
*Archaeolepidotus leonardi* Accordi, 1955, holotype, from the Monte Pic, Dolomites, stored in the Museum Gherdëina. [planned for page width]

Because a similar skeletal element in other taxa could not be found in the literature, it is not possible to assign PZO 16555 at low taxonomic rank, leaving it as Actinopterygii gen. et sp. indet.

> Class REPTILIA Laurenti, 1768
>
> [by A. S. Wolniewicz]
>
> Clade DIAPSIDA Osborn, 1903
>
> Clade SAUROPTERYGOMORPHA Wolniewicz et al., 2023
>
> Family SAUROSPHARGIDAE Li et al., 2011
>
> SAUROSPHARGIDAE indet.
>
> Figure 9

**FIGURE 9.**
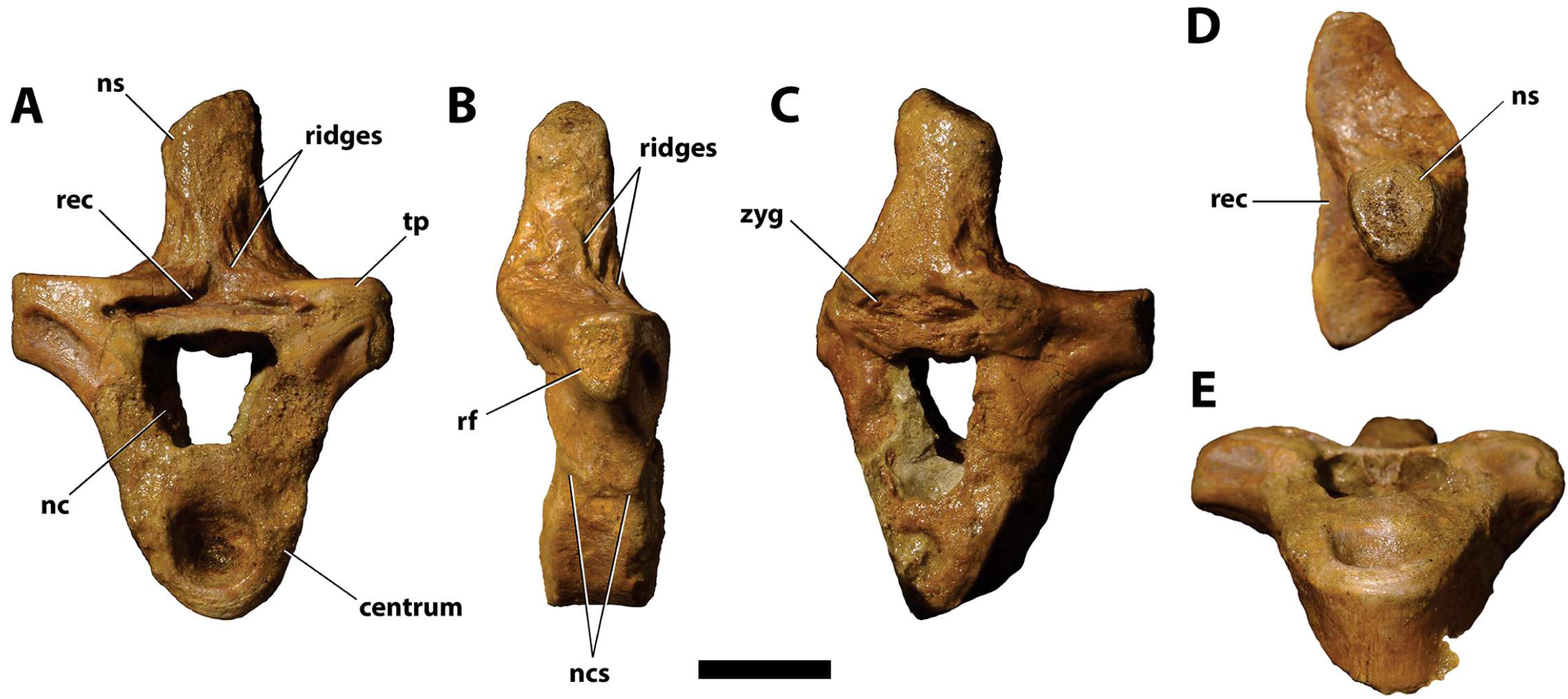
Saurosphargidae indet., PZO 773, an anterior postaxial cervical vertebra in **A**, anterior; **B**, right lateral; **C**, right posterolateral; **D**, dorsal; **E**, anteroventral views. Note that the posterolateral portion of the centrum remains embedded in matrix and is therefore not visible in **C** and **E**. Abbreviations: **nc**, neural canal; **ns**, neural spine; **ncs**, neurocentral suture; **rf**, rib facet; **rec**, recess ; **tp**, transverse process; **zyg**, zygantrum. Scale bar = 10 mm. [planned for page width]

**Referred specimen–** PZO 773, an isolated anterior postaxial cervical vertebra.

**Horizon, locality, and age–** Val Badia Member of the Werfen Formation, lower Spathian (late Early Triassic); western slope of the Weißhorn above the Bletterbach gorge.

**Description–** PZO 773 is identified as an anterior postaxial cervical vertebra of an indeterminate saurosphargid. The vertebra is extensively abraded, particularly on the anterodorsal surface of the centrum, the anterior surfaces of the neural arch pedicles, the left transverse process and the neural spine. Abrasion obscures details of surface morphology and has resulted in the loss of several anatomical features, including the zygapophyses, zygosphene and apex of the neural spine.

The centrum is deeply amphicoelous, as in *Sinosaurosphargis yunguiensis* and *Largocephalosaurus qianensis* (Li et al. 2011, 2014). It is relatively wide mediolaterally, reaching its greatest width dorsally, and is anteroposteriorly shorter than dorsoventrally tall, resembling the morphology of cervical vertebrae 3–5 in the holotype of *Largocephalosaurus qianensis* (Li et al. 2014). The lateral surfaces of the centrum face ventrolaterally and are slightly concave anteroposteriorly, giving the centrum a constricted appearance in ventral view. The ventral surface is transversely convex and lacks a ventral keel. In all these features, the centrum of PZO 773 closely resembles the postaxial cervical and anterior dorsal centra of other saurosphargids (Li et al. 2011, 2014).

The centrum and neural arch are co-ossified, although the neurocentral suture remains open and clearly visible on both sides of the vertebra. The neural arch pedicles are relatively tall dorsoventrally and wide mediolaterally, and they extend dorsolaterally from the centrum, like in *Sinosaurosphargis* and *Largocephalosaurus* (Li et al. 2011, 2014). In anterior view, the neural canal is approximately trapezoidal in outline, and is slightly wider mediolaterally than tall dorsoventrally, reaching its greatest width dorsally. In *Largocephalosaurus qianensis*, the neural canal in a posterior cervical vertebra is approximately as wide as tall (Li et al. 2014), whereas in *Sinosaurosphargis*, the neural canal of an anterior dorsal vertebra is markedly wider than tall (Li et al. 2011). The lateral walls of the neural canal are inclined posteromedially, such that in posterior view the neural canal appears proportionally much taller than wide.

The prezygapophyses and postzygapophyses are not preserved due to extensive abrasion of the vertebra. A mediolaterally wide, anteroposteriorly short and anteriorly incomplete zygosphene was reported in a posterior cervical vertebra of *Largocephalosaurus qianensis* (Li et al. 2014). A similar zygosphene appears to be present in PZO 773, but it is abraded. A mediolaterally wide and dorsoventrally narrow recess is located posterodorsal to the zygosphene. In a complete vertebra, this recess would have been laterally bordered by the prezygapophyses, as evidenced by a similar but shallower recess present between the prezygapophyses in an anterior dorsal vertebra of *Sinosaurosphargis* (Li et al. 2011). Posterodorsal to the neural canal, the neural arch forms a mediolaterally wide, posteroventrally inclined and slightly convex surface occupied by a shallow, mediolaterally wide, oval-shaped concavity representing the zygantrum. A nearly identical morphology of the posterior surface of the neural arch was reported in a disarticulated, anterior postaxial cervical vertebra in the referred specimen of *Largocephalosaurus qianensis* (Li et al. 2014).

The left transverse process is extensively abraded, but the right one is well-preserved. It is relatively tall dorsoventrally and short mediolaterally, like the transverse processes of a disarticulated postaxial cervical vertebra associated with the skull of the referred specimen of *Largocephalosaurus qianensis* (Li et al. 2014). A rib facet is visible on the lateral surface of the right transverse process; it is taller (dorsoventrally) than long (anteroposteriorly) and tapers ventrally. A distinct oblique ridge extends from the dorsolateral to the ventromedial part of the anterior surface of the transverse process, separating its dorsal and ventral portions and forming a noticeable “step”, such that the ventral portion is slightly depressed relative to the dorsal one. The posterior surface of the transverse process exhibits a shallow concavity near its base.

The surfaces of the neural spine are heavily abraded. The neural spine is robust and oval in cross-section, being wider mediolaterally than long anteroposteriorly. As preserved, the neural spine is dorsoventrally taller than the centrum, contrasting with the condition in all other saurosphargids, in which the neural spines of the cervical and dorsal vertebrae are shorter than the centrum (Li et al. 2011, 2014; Klein and Scheyer 2024). Arcuate, subvertical ridges are present on the anterolateral surfaces of the neural arch and increase in size from medial to lateral. Five ridges are preserved on the left side and three on the right. Similar ridges appear to ornament the anterolateral surfaces of a dorsal neural spine in a disarticulated posterior cervical vertebra in the referred specimen of *Largocephalosaurus qianensis* (Li et al. 2014).

In general morphology, PZO 773 closely resembles a disarticulated anterior postaxial cervical vertebra preserved in the referred specimen of *Largocephalosaurus qianensis* (Li et al. 2014), although its neural spine is proportionally taller. Although PZO 773 exhibits an autapomorphic, proportionally tall neural spine, we refrain from naming a new taxon due to the fragmentary nature of the material.

### Measurements

Total height (dorsoventral): 44.54 mm

Total width (mediolateral): 30.95 mm

Centrum length (anteroposterior): 10.14 mm

Centrum width (mediolateral, along dorsal surface): 14.95 mm

Centrum height (dorsoventral, anterior view): 13.83 mm

Neural canal width (mediolateral, along dorsal margin, anterior view): 10.50 mm

Neural canal height (dorsoventral, anterior view): 8.69 mm

Neural spine height (dorsoventral, incomplete): 18.15 mm

**Remarks–** Saurosphargidae is a small group of marine reptiles ranging from the Early to Middle Triassic of eastern and western Tethys, although fragmentary fossil material suggests that they may have survived into the Late Triassic (latest Norian–Rhaetian), at least in the western Tethys (Scheyer et al. 2022). Saurosphargids are currently represented by five taxa: *Prosaurosphargis yingzishanensis* from the Early Triassic (Olenekian) of Hubei Province, China (Wolniewicz et al. 2023); *Saurosphargis volzi* from the Middle Triassic (Anisian) of Poland and Germany (Huene 1936; Nosotti & Rieppel 2003; Klein & Scheyer 2024); and *Sinosaurosphargis yunguiensis*, *Largocephalosaurus polycarpon* and *Largocephalosaurus qianensis*, all from the Middle Triassic (Anisian) of Yunnan and Guizhou provinces, China (Li et al. 2011, 2014). Saurosphargids are characterised by several diagnostic morphological features, including a dorsal ‘rib-basket’, dorsal osteoderms, and broadened lateral gastral elements (Li et al. 2014; Wolniewicz et al. 2023). They were closely related to, or possibly even nested within, Sauropterygia. Some phylogenetic analyses have recovered saurosphargids as the sister group to Sauropterygia (e.g. Li et al. 2014; Wang et al. 2022), whereas others have recovered them as the sister group to Eosauropterygia (e.g. Simões et al. 2022; Wolniewicz et al. 2023).

PZO 773 represents an abraded anterior postaxial cervical vertebra and is therefore too fragmentary to justify the erection of a new saurosphargid taxon. Its morphology most closely resembles that of a disarticulated postaxial cervical vertebra associated with the skull in the referred specimen of *Largocephalosaurus qianensis* (Li et al. 2014), but the neural spine is proportionally much taller than in any currently known saurosphargid (Li et al. 2011, 2014; Klein and Scheyer 2024). However, the three-dimensional morphology of saurosphargid postaxial cervicals remains poorly documented in general, and is entirely unknown in *Sinosaurosphargis* and *Prosaurosphargis* (Li et al. 2011; Wolniewicz et al. 2023), limiting meaningful comparisons between taxa. Thus, while it is plausible that PZO 773 represents a new saurosphargid taxon, additional fossil material and more detailed studies of saurosphargid cervical morphology are required to confirm this.

PZO 773 represents one of only two unambiguous records of Saurosphargidae from the Early Triassic, the other being *Prosaurosphargis yingzishanensis* from the latest Spathian (Olenekian) of the Jialingjiang Formation, Hubei Province, southern China (Wolniewicz et al. 2023). *Saurosphargis volzi*, a saurosphargid known from the Gogolin Beds of Poland and the lower Muschelkalk of Schleusingen, Germany, is generally regarded as Anisian (Middle Triassic) in age (Huene 1936; Rieppel & Hagdorn 1997; Klein & Scheyer 2024). However, the lowermost part of the Gogolin beds is considered to be Early Triassic (latest Spathian, Olenekian) in age (Nawrocki and Szulc 2000), and eosauropterygian remains have been reported from these horizons (Kowal-Linka & Bodzioch 2012, 2017). Consequently, the possibility that *Saurosphargis volzi* extended into the Early Triassic cannot be completely ruled out, as the exact stratigraphic horizon of the type specimen from the Gogolin beds remains unknown. Unfortunately, the type specimen of *Saurosphargis volzi* was destroyed during World War II, making its precise stratigraphic provenance impossible to confirm.

PZO 773 represents one of the stratigraphically earliest known occurrences of Sauropterygomorpha—the clade that includes sauropterygians, saurosphargids, and their close relatives (Wolniewicz et al. 2023). Other early sauropterygomorph occurrences are represented by the eosauropterygian *Majiashanosaurus discocoracoidis* from the middle Spathian (radiometric age: 248.7 ± 1.0 Ma) Chaohu Fauna of the Nanlinghu Formation, Anhui Province, China (Jiang et al. 2014; Wang et al. 2025), and the saurosphargid *Prosaurosphargis* and the eosauropterygians *Hanosaurus hupehensis*, *Pomolispondylus biani*, and *Chusaurus xiangensis* from the latest Spathian (radiometric age: 246.7 ± 2.0 Ma) Nanzhang-Yuan’an Fauna of the Jialingjiang Formation, Hubei Province, China (Cheng et al. 2022; Wang et al. 2022; Liu et al. 2023; Wolniewicz et al. 2023). Sauropterygians have also been reported from the Spathian of the Sulphur Mountain Formation of British Columbia, Canada (Scheyer et al. 2019), as well as from the latest Spathian–earliest Anisian Gogolin Beds of Poland (Kowal-Linka & Bodzioch 2012, 2017) and the Alcova Limestone of Wyoming, USA (Storrs 1991; Lovelace & Doebbert 2015). However, precise radiometric ages are not available for these horizons. PZO 773 derives from the Val Badia Member of the Werfen Formation, which is regarded as early Spathian in age based on ammonoid biostratigraphy (Broglio Loriga et al. 1990; Posenato 2019; Zhang et al. 2019). This suggests that the specimen possibly represents the stratigraphically earliest known occurrence of a sauropterygomorph reported to date. However, due to the lack of precise radiometric dating for the Val Badia Member, direct age comparison with the Nanzhang-Yuan’an and Chaohu faunas is currently not possible.

PZO 773 potentially represents the stratigraphically earliest saurosphargid discovered to date and provides important insights into the timing of sauropterygian origins. If saurosphargids are indeed the sister group to Sauropterygia (e.g. Li et al. 2014; Wang et al. 2022), their presence in the early Spathian would suggest that the divergence between Saurosphargidae and Sauropterygia occurred no later than the early Spathian, perhaps in the aftermath of the end-Smithian extinction. Alternatively, if saurosphargids are nested within Sauropterygia (e.g. Simões et al. 2022; Wolniewicz et al. 2023), their presence in the early Spathian indicates that sauropterygian diversification must have proceeded very rapidly during the earliest Spathian. Although no saurosphargid or sauropterygian fossils are currently known from earlier than the early Spathian, it cannot be completely ruled out that the origins of the group extend back into the Smithian or even the Permian, as has recently been proposed for ichthyosaurs (Kear et al. 2023). New fossils from the latest Permian and earliest Triassic are crucial for testing these competing hypotheses on the timing of Mesozoic marine reptile origins.

The geographic origin of sauropterygomorphs remains debated. Traditionally, an eastern Tethyan origin has been proposed, as the stratigraphically earliest known sauropterygomorphs, represented by saurosphargids and eosauropterygians, have been recovered from middle–late Spathian (Olenekian, Early Triassic) fossil horizons of eastern Tethys (i.e., southern China) (Jiang et al. 2014; Li and Liu 2020; Wolniewicz et al. 2023; Wang et al. 2025). However, the discovery of PZO 773 challenges this view.

The specimen potentially represents the stratigraphically oldest known saurosphargid and originates from early Spathian strata of the western Tethys region (Italian Dolomites). The stratigraphic position of PZO 773 and its western Tethyan origin appear to support the hypothesis that saurosphargids—and possibly sauropterygomorphs more broadly—originated in the western Tethys no later than the early Spathian and dispersed eastward, reaching the eastern Tethys by the middle–late Spathian. A western Tethyan origin of Sauropterygomorpha is also corroborated by recent phylogenetic analyses, which recover several western Tethyan taxa as the earliest-diverging sauropterygomorphs. These include *Palatodonta bleekeri* (an early-diverging placodontiform or early-diverging sauropterygomorph) (Neenan et al. 2013; Wolniewicz et al. 2023), *Eusaurosphargis dalsassoi* (a potential sister taxon to Sauropterygia or early-diverging saurosphargid) (Nosotti and Rieppel 2003; Scheyer et al. 2017; Wang et al. 2022), and *Helveticosaurus zollingeri* (variably placed near or within Sauropterygomorpha) (Li et al. 2014; Wang et al. 2022; Wolniewicz et al. 2023), all known from the Middle Triassic of Europe. Notably, none of these early-diverging sauropterygomorphs (or cloesely related taxa) have been reported from the eastern Tethys, despite over 25 years of intensive fieldwork in Early to Late Triassic fossil deposits in southern China.

Numerous fragmentary remains of sauropterygians and likely related marine reptiles, that are potentially important for understanding the timing and geographic origins of Sauropterygomorpha, have been reported from the Early to Middle Triassic of Poland, Germany, and the Netherlands (e.g., Rieppel 1995; Surmik 2016; Spiekman & Klein 2021). While some of these fossils bear similarities to known taxa, they also exhibit morphological features that hinder confident assignment to specific sauropterygomorph groups. New discoveries of early-diverging sauropterygomorphs, detailed reinvestigation of available fossil specimens, and a more robust understanding of phylogenetic relationships among sauropterygians and their relatives are critical for resolving questions about the timing and geographic origins of this group.

### Radiolarians from rock sample with Ptycholepidae scales (PZO 16554) at the Seres section

The rock sample with Ptycholepidae scales (PZO 16554), collected from the surface at the Seres section, is a cherty limestone (wackestone) with abundant ostracods and radiolarians (Figure 10). The sample contains a rich, relatively well-preserved and diverse radiolarian fauna (Figure 11). The species composition of this assemblage reflects the typical radiolarian fauna of Buchenstein beds from localities in the southern Alps, Julian Alps and Transdanubian Central Range, as described by Dumitrica (1978a, b), Kozur and Mostler (1979, 1981, 1994), Dumitrica et al. (1980), Lahm (1984), and Ozsvárt et al. (2023). Forty-nine species could be identified from the fauna, of which 61% were Spumellaria, 26% Nassellaria, and 13% Entactinaria. Therefore, the diversity of spumellarians and nassellarians significantly outnumber entactinarians by a ratio of about six to one (Table 1).

**FIGURE 10.**
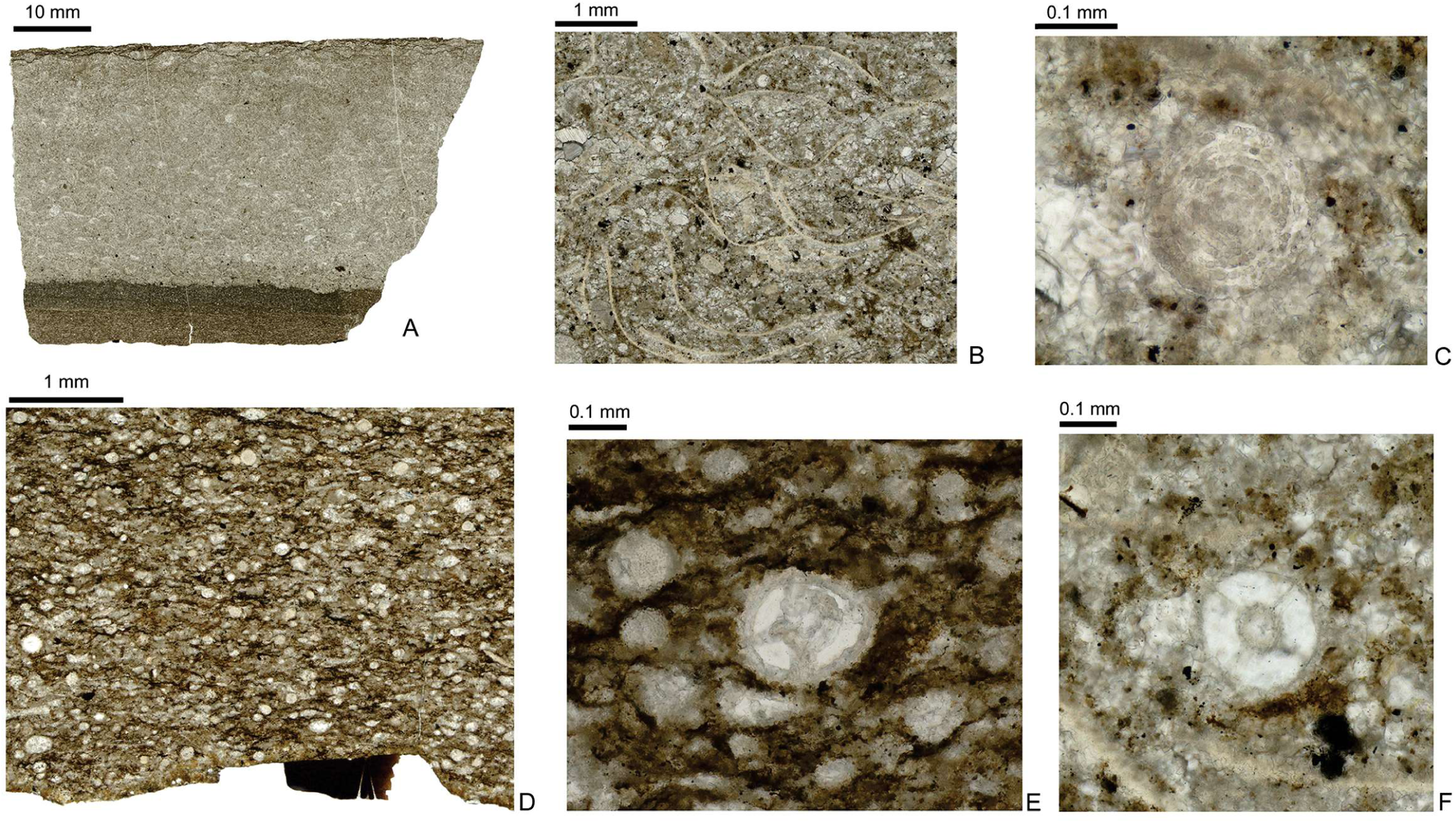
Thin section prepared from the rock slab with Ptycholepidae scales (PZO 16554) in Figure 6, Anisian–Ladinian (Middle Triassic) Buchenstein Formation (Illyrian according to radiolarians, latest Olenekian–earliest Anisian according to U–Pb dating). **A**, Thin section image, showing dark layer where the scales were found on lower surface, and light-colored layer with abundant ostracods; **B**, close-up of ostracods in light-colored layer, **C**, close-up of a radiolaria (Oertlispongidae) in the light-colored layer; **D**, dark-colored layer with a scale (dark brown) in cross section in lower surface; **E**, close-up of a radiolaria (*Lahmosphaera*) in the dark-colored layer likely; **F**, close-up of a radiolaria (*Lahmosphaera*) in the light-colored layer. [planned for page width]

**FIGURE 11.**
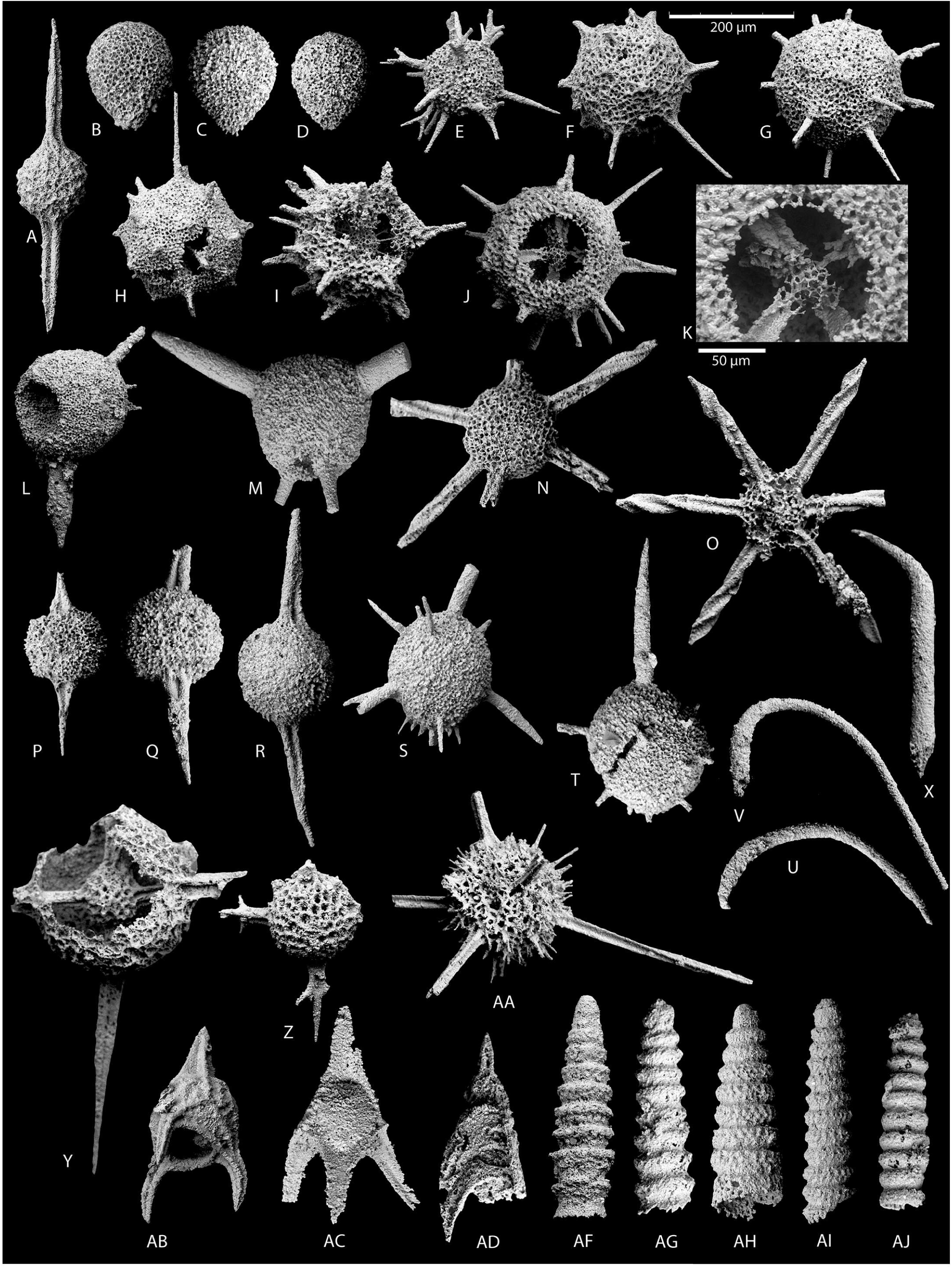
Radiolarians found in the rock slab with Ptycholepidae scales and other elements (PZO 16554), Anisian–Ladinian (Middle Triassic) Buchenstein Formation (Illyrian according to radiolarians). **A**, *Pseudostylosphaera longispinosa* Kozur and Mostler, 1981; **B–D**, *Glomeropyle* cf. *aurora* Aita, 1999; **E**, *Katorella trifurcata* Kozur and Mostler, 1981; **F,** *Triassospongosphaera austriaca* (Kozur and Mostler, 1979); **G,** *Triassospongosphaera triassica* (Kozur and Mostler, 1979); **H,** *Triassospongosphaera multispinosa* (Kozur and Mostler, 1979); **I,** *Astrocentrus latispinosus* (Kozur and Mostler, 1979); **J–K,** *Astrocentrus*? cf. *pulcher* Kozur and Mostler, 1979, **K**: initial spicule system; **L,** *Neopaurinella* cf. *tumidaspina* Kozur and Mostler, 1994; **M,** *Neopaurinella* cf. *ladinica* Kozur and Mostler, 1994; **N–O**, *Relindella ruesti* (Kozur and Mostler, 1981); **P–Q**, *Spongotortilispinus koppi* (Lahm, 1984); **R,** *Spongopallium* sp.; **S,** *Paroertlispongus kozuri* Ozsvárt, 2023; **T,** *Paroertlispongus multispinosus* Kozur and Mostler, 1981; **U–V** *Oertlispongus inaequispinosus* Dumitrica et al., 1980; **X,** *Paroertlispongus weddigei* (Lahm, 1984) **Y,** *Lahmospharea* sp.; **Z,** *Lahmospharea trispinosa* (Kozur and Mostler, 1979); **AA,** *Ticinosphaera mesotriassica* (Kozur and Mostler, 1981); **AB,** *Hozmadia costata* Kozur and Mostler, 1994; **AC,** *Eonapora mesotriassica* Kozur and Mostler, 1981; **AD,** *Hinedorcus alatus* Dumitrica et al., 1980; **AF,** *Triassocampe scalaris* Dumitrica et al., 1980; **AG,** *Paratriassocampe gaetanii* Kozur and Mostler, 1994; **AH,** *Pararuesticyrtium eofassanicum* Kozur and Mostler, 1994; **AI,** *Annulotriassocampe campanilis campanilis* Kozur and Mostler, 1994; **AJ,** *Annulotriassocampe campanilis longiporata* Kozur and Mostler, 1994. [planned for page width]

**TABLE 1.** The list of radiolarian taxa found in the rock sample with Ptycholepidae scales (PZO 16554) at the Seres section, and their known ranges (from Ozsvárt et al. 2023).

The most dominant spumellarians are spongy forms such as from the Intermediellidae group (*Paurinella, Neopaurinella, Triassospongosphaera, Katorella* and *Astrocentrus*) and the Oertlispongidae (*Paroertlispongus, Oertlispongus*). Additionally, important and abundant elements of the radiolarian fauna include some precariously classified forms such as *Lahmosphaera* and *Ticinosphaera* (Table 1). Nearly a quarter of the radiolarian fauna are nassellarians, predominantly monocyrtids such as *Hozmadia*, and *Eonapora*, although the multicyrtid Ruesticyrtiidae (*Pararuesticyrtium, Triassocampe* and *Annulotriassocampe*) are also important. The entactinarians are dominated by Hindeosphaerinae: especially *Pseudostylosphaera* and Parentactiniidae such as *Glomeropyle* (Table 1). Interestingly, specimens of the genus *Glomeropyle* Aita & Bragin have so far been known mainly from areas outside the Tethyan realm, such as New Zealand, or from boreal areas such as Siberia. For this reason, the occurrence of *Glomeropyle* is completely unique in the Buchensten Basin.

### Age of the rock sample with Ptycholepidae scales (PZO 16554)

The composition of the radiolarian fauna from this rock sample (PZO 16554) is 95% identical to the radiolarian fauna of the Frötschbach and Seceda sections published from the Dolomites (Ozsvárt et al., 2023). Hence, the most probable age of this rock sample is late Illyrian (late Anisian), as this radiolarian assemblage belongs to the middle part of the *Spongosilicarmiger italicus* Radiolaria Zone from the Buchenstein Formation. This horizon is equivalent in age to the *Reitziites reitzi* Ammonoid Zone (*Reitzi* Subzone to *Avisianum* Subzone) and *Paragondolella trammeri* Conodont Zone.

U–Pb LA–ICPMS data from the Ptycholepidae fish scales and carbonate host rock are presented in Supplementary Material (Table S1). The fish scales show U content of about 3 ppm and Pb content of 0.3 ppm, whereas the U content in the limestone is about 1 ppm and Pb content is 2 ppm. Both scales and host rock show null ^232^Th content.

The host rock yields an age of 248 ± 1 Ma with a PbC of 0.835 ± 0.002, showing a good spread on the U/Pb ratio along the mixing line (Figure 12). The PbC value of 0.835 is consistent with the average crustal model-based common Pb for this age (∼0.85, Stacey & Kramers, 1975). In contrast, the scales yield a cluster of data below the concordia curve at about 600 Ma (Figure 12).

**FIGURE 12.**
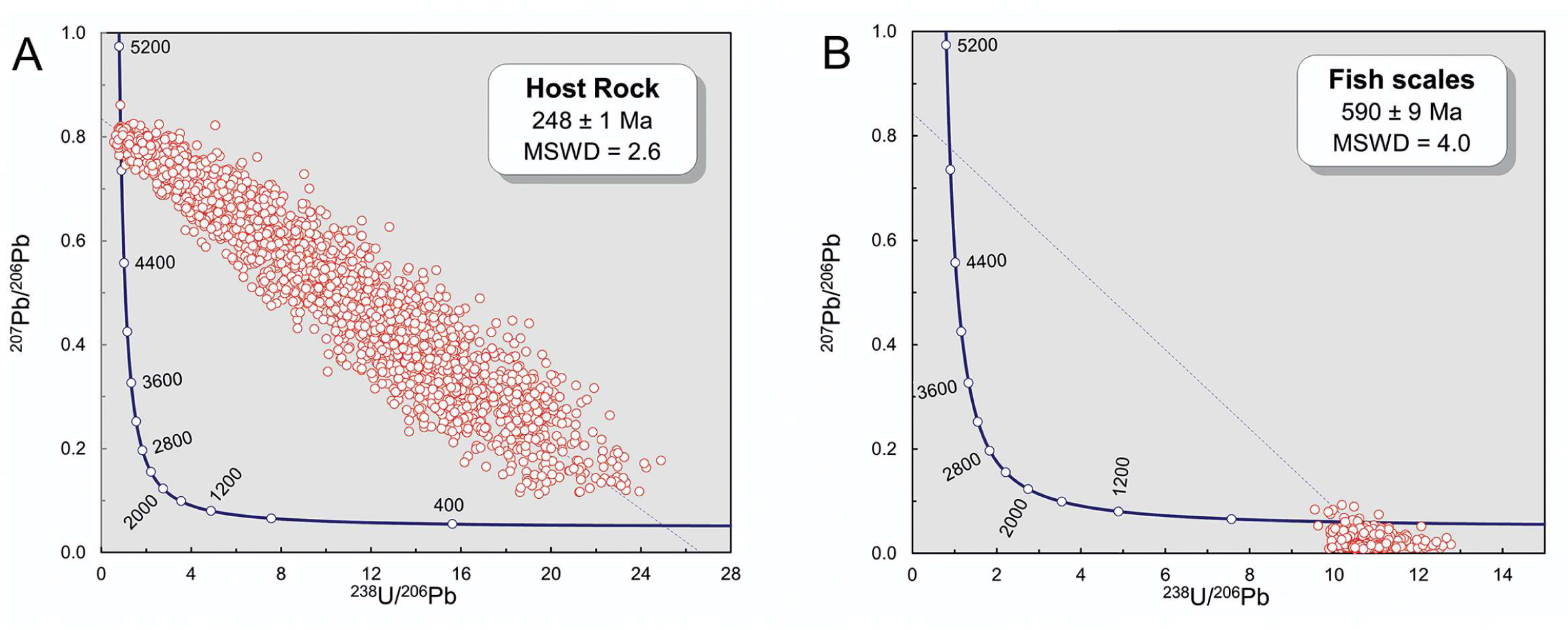
**A**, U–Pb dating of rock slab with Ptycholepidae scales and other elements (PZO 16554) indicating an age of 248 ± 1 Ma (latest Olenekian–earliest Anisian). **B**, U–Pb dating of scales on the sample PZO 16554, indicating the age of 590 ± 9 Ma (Ediacaran). [planned for page width]

## DISCUSSION

Using both surface and bulk sampling methods has been shown to capture diversity more accurately in marine invertebrates, as each method covers different size ranges that do not fully overlap (e.g., Forcino & Stafford 2020; Karapunar et al. 2022). Using these two sampling methodologies helped to recover several vertebrate fragments and revealed the youngest known Paleozoic occurrence of *Bobasatrania,* just below the end-Permian extinction horizon (given that the horizon of the Changhsingian material from Greenland was not specified; Stemmerik et al. 2001; Böttcher 2014), the presence of *Acrodus* in the shallow shelf environment, and one of the oldest and first western Tethyan occurrences of Saurosphargidae in the shallow shelf setting in the Spathian of the Dolomites.

The host rock could be accurately identified in most of the surface samples due to the outcrop not being a scree deposit and the presence of characteristic lithological features or fossil content. However, the rock sample with Ptycholepidae scales (PZO 16554) was further investigated by making thin sections and performing U–Pb dating to assign it to a horizon in the section, as it was collected from a scree deposit. Unexpectedly, the thin sections revealed the presence of Radiolaria, which have not been reported from either the Bellerophon or Werfen Formation, which is consistent with the global phenomenon known as the “chert gap” (e.g., Beauchamp & Baud 2002; Yang et al. 2022; Foster et al. 2023). The rock sample with scales was found to be of Illyrian (latest Anisian) age according to the radiolarians, even though it was found in scree dominated by rocks from the Bellerophon and Werfen formations, and the youngest part of the succession in the section is the Griesbachian Mazzin Member, which is overlain uncomformably by the Illyrian Richthofen Conglomerate with rounded pebbles and cobbles (Vachard & Krainer 2022). U–Pb dating of the host rock gives an age of 248 ± 1 Ma (Olenekian–Anisian age), slightly differing from the age indicated by the radiolarian biostratigraphy.

The U–Pb results on the scales cannot be interpreted in terms of age, partly because the apparent age would be much too old and because the data cluster beneath the concordia curve (Figure 12). The most reasonable explanation is that the ganoine-bearing actinopterygian scales were subjected to ^206^Pb enrichment during diagenesis (Figure 12). Fink et al. (2024) reported an unexpectedly young age in ganoine-bearing scales of gars (Lepisosteidae) from the Cretaceous based on (U-Th)/He dating, and suggested that the age discrepancy was caused by diagenetic alteration. The ^206^Pb enrichment mechanism in the studied actinopterygian scales is most likely due to preferential concentration of ^226^Ra, which is a ^238^U series radionuclide with a half-life of about 1,600 Ma. It is present in seawater from the decay of dissolved U, but it would require an enormous concentration factor to explain the amount of ^206^Pb found in the sample. The presence of this ^206^Pb suggests that during diagenesis the scales contained about 3 ppm of ^226^Ra, assuming that diagenesis lasted for a period shorter than the half-life. This would have produced a radiation level of about 11 disintegrations per second in a milligram of fossil material. The ^206^Pb enrichment mechanism in the studied actinopterygian scales is unknown and deserves further investigation, perhaps incorporating other phosphatic fossils from age constrained horizons.

Coincidentally, similar to the rock sample with ptycholepid scales found in a scree deposit dominated by Werfen Formation rocks at the Permian–Triassic outcrop (this study), *Archaeolepidotus leonardii* Accordi, 1955, was also collected from landslide material predominantly composed of Werfen Formation rocks (Accordi 1955, p. 6). Hence, Accordi (1955) presumed that the rock sample yielding *Archaeolepidotus leonardii* originated from the lower Triassic Siusi Member, although he described the color of the rock as dark grey (“grigio-scuro”; Accordi 1955, p. 6), suggesting the Bellerophon Formation. In our specimen, radiometric dating and radiolarian biostratigraphy suggest that the scale-bearing rock specimen originated from the Anisian. This finding underlines the unreliability of using lithological characteristics alone to determine the age of scree deposits. The specimens found by Accordi (1955; *Bobasatrania ladina* and *Archaeolepidotus leonardii*) might have come from the Changhsingian deposits (Renato Posenato, pers. comm. in Brinkmann et al. 2010). According to Accordi (1955, p. 21), *Bobasatrania ladina* was found in a gray, marly layer teeming with *Myacites* [=*Unionites*], located two to three meters above *Gymnocodium*-bearing bituminous rocks. *Unionites* range from the Changhsingian Bellerophon Formation to the Early Triassic Werfen Formation in the Dolomites (Prinoth & Posenato 2023), and *Gymnocodium* occurs both in the Bellerophon Formation and in the oolitic beds of the Tesero Member (Vachard & Krainer 2022). The color of the deposits and high abundance of *Unionites* and *Gymnocodium* suggest that Accordi’s material of *B. ladina* likely came from the Changhsingian Bellerophon Formation. To confirm the horizon of origin, the host rock should be restudied for its fossil content.

The isolated teeth of *Bobasatrania* from the uppermost Bellerophon Formation at Seres and Siusi in bulk samples, and the tooth plate in the surface-collected sample from the Bletterbach section, corroborate the existence of this taxon in the Changhsingian. They also show that its geographic range extended from equatorial latitudes to mid-latitudes before the extinction (Accordi 1955; Stemmerik et al. 2001; this study). *Bobasatrania* is one of the few fish genera that survived the Permian–Triassic mass extinction, albeit without expanding into new habitats (Tintori et al. 2014; Romano et al. 2016). Geographic range is the strongest correlate with survival likelihood, though this relationship weakens or disappears during mass extinction events (Payne & Finnegan 2007; Payne & Clapham 2011; Finnegan et al. 2023). The wide geographic distribution of *Bobasatrania* across latitudes possibly indicates a wider range of environmental tolerance, such as for temperature. Thermal tolerance range is an important determinant of species distribution in modern oceans (e.g., Day et al. 2018) and a good predictor of extinction likelihood for marine genera in the past (Reddin et al. 2022). The geographic range of *Bobasatrania*, extending from tropical paleolatitudes (Dolomites) to mid-paleolatitudes (Greenland), might have increased the chance of species survival, and contributed to the global distribution of the genus in the aftermath of the mass extinction event.

Specimens identified as *Acrodus*, with or without certainty, were previously reported from Lopingian (upper Permian) deposits in tropical paleolatitudes of the Paleo-Tethys (Italy: Brandt 2021; Hungary: Mihály & Solt 1981; Iran: Hampe et al. 2013) and the Zechstein Basin (Latvia: Dankina et al. 2020), along with possible occurrences in lower to middle Permian deposits in tropical paleolatitudes of the Panthalassa (Cisuralian, Texas, USA: Johnson 1981; Ginter et al. 2010; Roadian [Guadalupian], Arizona, USA, Hodnett et al. 2011). The *Acrodus* specimen studied herein is different in morphology compared to previously reported occurrences and corroborates the presence of this genus in the Changhsingian at mid-latitudes prior to the extinction event. In the Early Triassic, *Acrodus* was only reported from mid to high paleolatitudinal settings, such as Pakistan (Waagen 1895), Madagascar (Thompson 1982), Svalbard (Błażejowski 2004; Bratvold 2018), south Primorye, Russia (Yamagishi in Shigeta et al. 2009), and possibly Kashmir, India (Sahni & Chhabra, 1976). The apparent range shift of the genus towards higher paleolatitudes across the Permian/Triassic boundary is congruent with the expected range shift during the hyperthermal coinciding with the mass extinction event (up to 12°C warming across the P/Tr boundary; Gliwa et al. 2022) and the previously shown polewards shift in diversity gradient of bony fishes (Romano et al. 2016, fig. 9A). A global compilation of shark genera also shows selected disappearance of chondrichthyans in tropical paleolatitudes across the Permian–Triassic transition (Koot 2013; figs 8.29–8.31). Further studies may clarify whether this apparent range-shift pattern in fishes is a genuine signal or results from i) regional or temporal taphonomic differences due to environmental conditions (e.g., dysoxia in some basins such as Svalbard and Greenland; Twitchett 1999), ii) species-area relationships (e.g., higher extensive coastlines in mid latitudes than tropical latitudes), or iii) lack of sampling effort or outcrop preservation.

Although fully aquatic reptiles (Sauropsida) were present as a minor component in the Permian (e.g., *Mesosaurus*), several new marine reptile groups appeared as early as the Spathian in post-extinction marine communities (Ichthyosauromorpha and Sauropterygomorpha, the latter represented by Saurosphargidae and Eosauropterygia; Scheyer et al. 2019; Kear et al. 2023; 2024; Wang et al. 2022; Wolniewicz et al. 2023; Wang et al. 2025). The Triassic witnessed the explosive radiation of these and other marine reptile groups, and their eventual occupancy of the highest trophic levels in aquatic ecosystems during the Mesozoic (e.g., Jiang et al. 2023). The newly described saurosphargid vertebra (PZO 773) represents one of the earliest records of this group worldwide and the first unambiguous occurrence of the group from the western Tethyan Early Triassic. Apparently, saurosphargids dispersed from west to east in the Tethys by the latest Spathian (Wolniewicz et al. 2023) and were one of the earliest top predators in newly restructured Mesozoic marine communities of the Early Triassic.

The saurosphargid specimen (PZO 773) represents the first marine reptile fossil discovered in the Lower Triassic Werfen Formation of the Dolomites, after nearly two hundred years of collection and research. Up until now, Early Triassic vertebrates in the Werfen Formation in the Dolomites (Italy) were known only from tiny conodonts, possibly feeding on zooplankton, and perhaps fish, suggested by the trace fossil *Undichna* from the Smithian (Ronchi et al. 2018). In the Spathian deposits of the Werfen Formation in the Northern Calcareous Alps, Austria, vertebrate remains have been limited to only a few actinopterygians (*Gyrolepis*?, *Saurichthys* and *Colobodus*; Mostler & Rosner 1984). As discussed above, the Early Triassic age assignments for *Bobasatrania* and *Archaeolepidotus* still require confirmation.

The new saurosphargid record from the early Spathian marks the appearance of a new ecological niche, a relatively large-bodied reptilian predator, that was not present in the Changhsingian Bellerophon Formation. In the Changhsingian, the apex predator in the Dolomites community was perhaps the chondrichthyan *Acrodus* sp. reported herein, although it occupied a different, likely durophagous niche. The Spathian interval in the Werfen Formation also marks the reappearance of cephalopods (Broglio Loriga et al. 1990), following a time gap in their record since their disappearance in the Changhsingian (e.g., Prinoth & Posenato 2007). The saurosphargid can be considered the apex predator in the Spathian Dolomite community, as no other predator of comparable size has been reported from the Early Triassic of the Dolomites to date. Its presence suggests that this apex predator niche emerged before the full recovery of the benthos in the Dolomite community (e.g., see Hofmann et al. 2015; Friesenbichler et al. 2021), and argues against stepwise, bottom-up recovery in trophic webs postulated by Chen & Benton (2012) and previously also refuted by Scheyer et al. (2014).

Scheyer et al. (2014) proposed that the Early Triassic diversity of large marine vertebrates may be underestimated, as early marine reptiles likely inhabited shallow waters before evolving a fully pelagic lifestyle (Motani and Vermeij 2021). Their fossilization potential in such environments would have been lower compared to that of marine reptiles preserved in pelagic sediments, due to the higher likelihood of mechanical disarticulation in shallow-marine settings. The presence of the earliest saurosphargid record in a shallow-marine setting, represented by a single vertebra from the Dolomites (this study), supports this view. Our finding highlights the need for increased research and fieldwork efforts in shallow-marine deposits to uncover the earliest representatives of marine reptiles in the Early Triassic.

Fieldwork and sampling practices are the groundwork for paleontological and geological data collection. Therefore, they need to be conducted carefully to minimize potential error in data production, such as age or formation assignments to collected fossils (e.g., Fürsich et al. 2023). Although an extreme example, a misassignment of Werfen deposits to the Bellerophon Formation and taxonomic misidentifications previously resulted in complete misassignment of all Early Triassic Werfen taxa to Paleozoic species (Airaghi 1907). As discussed above, there are also uncertainties regarding the age and horizon of *Bobasatrania ladina* and *Archaeolepidotus leonardii* (Accordi 1956). Taxonomic or age misassignments can heavily alter our inference of changes in biodiversity and community structure in the past. To minimize the errors, interdisciplinary studies involving taxonomy combined with stratigraphical, geochemical, and geochronological studies are needed.

## CONCLUSION

New surface and bulk sample collections from several Permian–Triassic sections in the Dolomites, along with reexamination of specimens from museum collections, revealed new and previously undescribed vertebrate fossils from the Bellerophon and Werfen formations: *Bobasatrania* aff. *ladina*, *Bobasatrania* sp., *Acrodus* sp. and Actinopterygii indet. from the Changhsingian, prior to the extinction event; and Saurosphargidae indet. from the lower Spathian of the Werfen Formation during the post-extinction recovery period. The findings show the disappearance of chondrichthyans and osteichthyans from the studied tropical shallow shelf setting in the western Paleo-Tethys during the extinction interval, potentially related to the global hyperthermal event, and the appearance of a new top predator niche before full recovery of ecological and taxonomic diversity among the benthos. These discoveries enhance our understanding of the community structure across the Permian–Triassic extinction event in the Dolomites. The new vertebrate fossils presented herein demonstrate that there is still much to discover after nearly two centuries of paleontological research in the Dolomites, both in the field and in museum collections.

## ACKNOWLEDGEMENTS

Andrea Tintori (University of Milano) is acknowledged for preparation of the saurosphargid vertebra. Paulina Moroder (Museum Gherdëina) and Andrea Tintori are thanked for providing the pictures of *Bobasatrania* and *Archaeolepidotus*. We thank Martina Bonetto (Museum of Nature South Tyrol) for accessioning the specimens and providing collection numbers, and Peter Daldos (Geomuseum Radein) for photographing the tooth plate. Imelda Hausmann (SNSB-BSPG) is thanked for her assistance during ASW’s visit to the Bavarian State Collections for Palaeontology and Geology to study the saurosphargid vertebra. ASW thanks Jun Liu (Hefei University of Technology) and Adam Rytel (Institute of Paleobiology, Polish Academy of Sciences) for insightful discussions on Early Triassic marine reptile faunas. BK, SZB, FG and WJF were funded by Deutsche Forschungsgemeinschaft (project no. FO1297/1-1). FG received further funding from the Alexander von Humboldt Foundation. ASW was funded by the Bekker Programme of the Polish National Agency for Academic Exchange (grant no BPN/BEK/2022/1/00194).

## AUTHOR CONTRIBUTIONS

BK, EK, WJF designed the project; BK, ASW, CR, PO, HR-B, EK, AL-A, MB gathered the data; BK, ASW, CR, PO, HR-B analyzed the data and drafted the manuscript. All authors edited the manuscript.

## DATA AVAILABILITY STATEMENT

The U–Pb LA–ICPMS data necessary to reproduce the results of the manuscript are available as a supplementary file.

## SUPPLEMENTARY FILE(S)

Supplementary_Material_Table_S1_U-Pb data.xlsx: U–Pb LA–ICPMS data from the Ptycholepidae fish scales and carbonate host rock.

